# PerturbNet predicts single-cell responses to unseen chemical and genetic perturbations

**DOI:** 10.1101/2022.07.20.500854

**Authors:** Hengshi Yu, Joshua D. Welch

## Abstract

Small molecule treatment and gene knockout or overexpression induce complex changes in the molecular states of cells, and the space of possible perturbations is too large to measure exhaustively. We present PerturbNet, a deep generative model for predicting the distribution of cell states induced by unseen chemical or genetic perturbations. Our key innovation is to use high-throughput perturbation response data such as Perturb-Seq to learn a continuous mapping between the space of possible perturbations and the space of possible cell states.

Using Sci-Plex and LINCS datasets, PerturbNet can accurately predict the distribution of gene expression changes induced by unseen small molecules given only their chemical structures. PerturbNet also accurately predicts gene expression changes induced by shRNA, CRISPRi, or CRISPRa perturbations using a perturbation network trained on gene functional annotations. Furthermore, self-supervised sequence embeddings allow PerturbNet to predict gene expression changes induced by missense mutations. We also use PerturbNet to attribute cell state shifts to specific perturbation features, including atoms and functional gene annotations. Finally, we leverage PerturbNet to design perturbations that achieve a desired cell state distribution. PerturbNet holds great promise for understanding perturbation responses and ultimately designing novel chemical and genetic interventions.

## Introduction

Recent experimental developments have enabled high-throughput single-cell molecular measurement of response to drug treatment. A high-throughput chemical screen experiment usually involves a large number of cells and multiple treatments, where each cell receives a kind of drug treatment and is impacted in a distinct manner [1, 2]. Understanding how drugs influence cellular responses helps discover treatments with desired effects, potentially benefiting a myriad of therapeutic applications.

Unlike chemical perturbations, whose direct gene targets are generally unknown, genetic perturbations are designed to directly knock out or activate one or multiple target genes. The activation or knockout of these genes will not only influence their own expression, but also impact other genes through a complex network of downstream gene regulatory interactions. The Clustered Regularly Interspaced Short Palindromic Repeats (CRISPR) technology allows precise design of genetic mutants through genome editing [3]. More recently, CRISPR has been combined with transcriptional activators (CRISPRa) or repressors (CRISPRi) tethered to a deactivated version of the Cas9 protein (dCas9) to enable activation or inhibition of target genes. The Perturb-seq technology combines CRISPR/Cas9 and single-cell RNA-sequencing (scRNA-seq) to measure single-cell responses to pooled CRISPR guide RNA libraries [4]. Perturb-seq measures cellular responses at single-cell resolution, revealing how cell states are impacted by genetic perturbations, and has been utilized for many biomedical applications [5, 6, 7, 8, 9]. However, because Perturb-seq experiments can only measure limited numbers of perturbation loci, it is not feasible to directly measure single-cell responses for each potential genetic perturbation.

Recent studies have shown that genetic perturbations can induce shifts in cell state, causing the cells to preferentially occupy certain cell states while disfavoring others. For example, [10] observed that CRISPR activation (CRISPRa) of distinct pairs of genes in K562 cells induced some cells to shift toward erythroid, granulocyte, or megakaryocyte-like states or arrest cell division. Inducing missense mutations in *KRAS* and *TP53* caused “a functional gradient of states” in A549 cells[8].

That is, in a particular tissue under homeostatic conditions, there is a wild-type distribution of cellular gene expression states *p*(***X***). Treating cells with a perturbation *G* changes their cell state distribution to some new *p*(***X***|*G*). Our goal is to predict these perturbation-specific distributions by developing deep generative models to sample from the cell state distribution for any perturbation.

Several recent methods have been developed for modeling single-cell perturbation effects. The variational autoencoder scGen predicted single-cell data from new combinations of treatment and cell type using latent space vector arithmetic [11]. Another method, Dr. VAE, also explored the dependency of the latent space on treatments [12]. For genetic perturbations, *Norman et al.* (2019) used Perturb-seq data to identify genetic interactions from paired gene knockouts [10]. *Lotfollahi et al.* (2020) proposed a conditional variational autoencoder (VAE) framework with representations under two treatment conditions [13] balanced using a similarity score of their counterfactual inference [14]. *Burkhardt et al.* (2021) identified perturbation effects over the cellular manifold using graph signal processing tools [15]. The compositional perturbation autoencoder framework [16] generates single-cell data under new combinations of observed perturbations using latent space vector arithmetic. *Yeo et al.* (2021) proposed a generative model using a diffusion process over a potential energy landscape to learn the underlying differentiation landscape from time-series scRNA-seq data and to predict cellular trajectories under perturbations [17]. Linear models were also used to estimate the impact of perturbations on high-dimensional scRNA-seq data [4] or infer gene regulatory interactions from perturbations [18].

The key limitation of existing approaches is that they focus only on predicting new combinations of treatments and/or cell types, and thus cannot predict the effects of a completely unseen perturbation. Additionally, many of the existing approaches treat perturbations as independent from cell state, making it impossible to accurately predict the effects of perturbations that specifically promote or disfavor certain cell states. To address these limitations, we propose PerturbNet, a novel and flexible framework that can sample from the distribution of cell states given only the features of a new perturbation. The PerturbNet model connects drug treatment or genetic perturbation information and cell states using a conditional normalizing flow [19], enabling translation between perturbation domain and single-cell domain [20]. The PerturbNet framework is generally applicable to any type of high-throughput measurement of drug treatments or genetic perturbations, such as those of scRNA-seq data. Importantly, PerturbNet makes distributional predictions for both observed and unseen treatments.

We show that PerturbNet can effectively predict the distribution of gene expression profiles induced by a variety of chemical and genetic perturbations. Using microarray data from the Library of Interconnected Network Signatures (LINCS) and scRNA-seq data from the Sci-Plex dataset, PerturbNet predicts the effects of small molecule treatment. Additionally, we show that PerturbNet can predict the effects of CRISPR activation and CRISPR-induced missense mutations from Perturb-seq data.

We further demonstrate that the predictive capability of PerturbNet is useful for two important downstream applications: (1) implicating key perturbation features that influence cell state distributions and (2) designing optimal perturbations that achieve desired effects. We interpret our predictive model to reveal key components or functions in a chemical or genetic perturbation that influence the cell state. We further develop an algorithm to design perturbations that optimally translate cells from a starting cell state to the desired cell state.

## Results

### PerturbNet maps perturbation representations to cell states

PerturbNet consists of three neural networks: a perturbation representation network, a cellular representation network, and a network that maps from perturbations to cell states (Fig. 1a). The representation networks are first trained separately to encode large numbers of perturbations and cell profiles into latent representations. Then the mapping network uses high-throughput perturbation response data such as Perturb-seq, in which both perturbation and cell states are observed, to learn a continuous mapping between the space of possible perturbations and the space of possible cell states. The intuition behind our approach is that perturbations and cells each have an underlying structure–that is, they lie in some low-dimensional space–and the effect that a perturbation exerts on cell state is given by some unknown function that maps between the two spaces (Fig. 1b). The mapping from perturbations to cell states is highly complex and not one-to-one; cells may exist in many states after a particular perturbation, and distinct perturbations may induce similar cell state distributions. To capture such complex relationships, we implement the mapping network with a conditional invertible neural network (cINN), which fits a conditional normalizing flow and can approximate arbitrary conditional distributions. Our approach is inspired by the idea of network-to-network translation, which has been used to generate images conditioned on text descriptions [21].

**Fig. 1.**
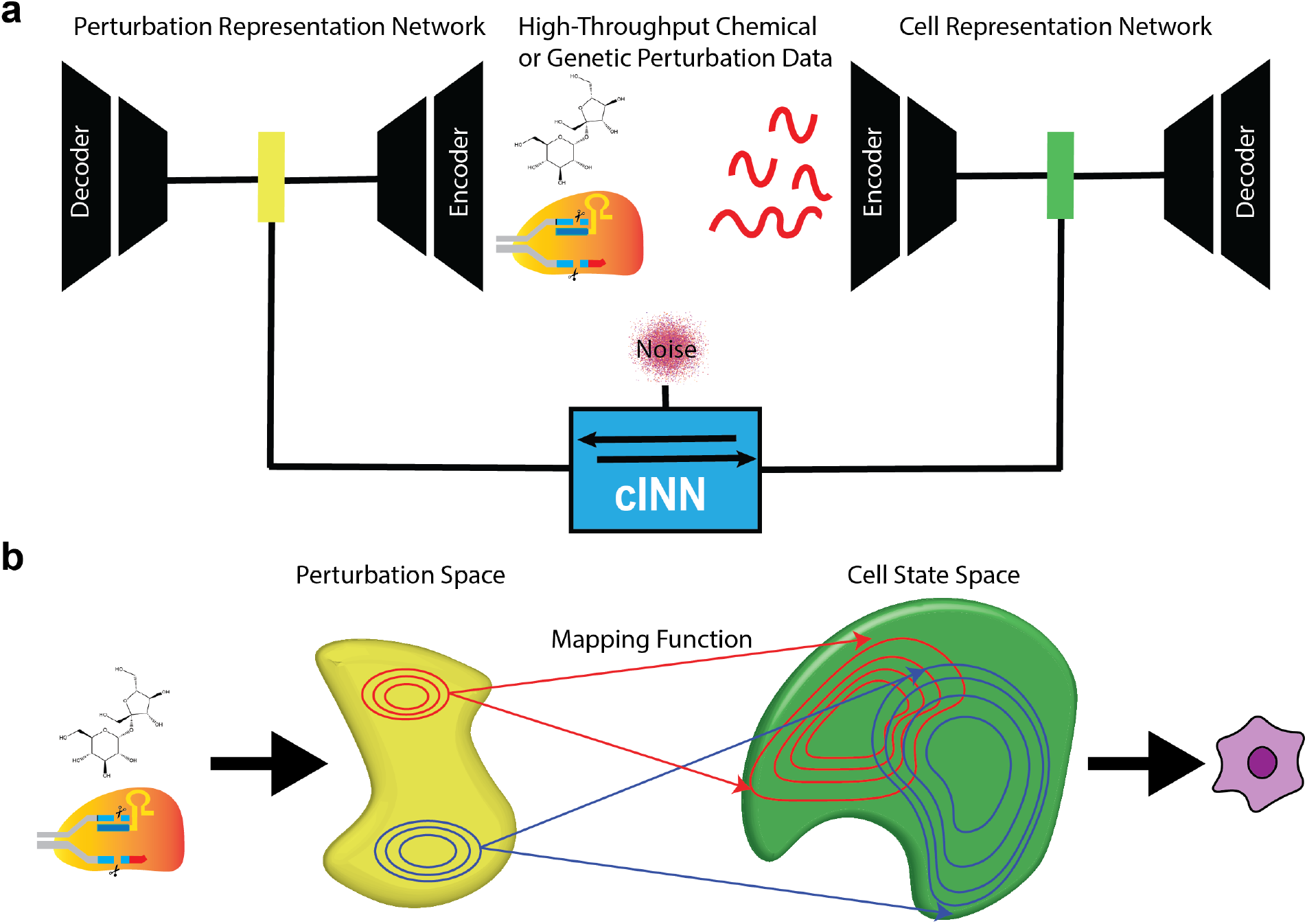
PerturbNet maps perturbation representations to cell states. **a** PerturbNet uses two neural networks separately trained to encode large numbers of chemical or genetic perturbations (left) and cell profiles (right) into latent spaces. A conditional invertible neural network (cINN) learns to map points in perturbation space to cell state space using high-throughput measurements of perturbation effect. **b** PerturbNet can then predict the gene expression changes induced by an unseen perturbation by encoding the perturbation, passing its representation through the cINN, and decoding the resulting cell states.

After training, PerturbNet can make predictions about the cell states induced by a new perturbation. To do this, a description of the perturbation–such as the chemical structure of a small molecule or the identities of the genes knocked out–is first encoded into the perturbation space. Then, this location in the perturbation space is fed into the mapping network, whose output gives the distribution of locations in the cell state space induced by the perturbation. These latent cell representations can subsequently be decoded into high-dimensional gene expression levels to predict the perturbation responses of individual genes.

The PerturbNet framework has several key advantages. First, the perturbation and cell representation networks are fully modular, allowing a variety of architectures to be used depending on the data type. For example, we can use convolutional and recurrent networks for representing small molecule structures, multilayer perceptrons for gene expression data, or transformer architectures for sequence data. A second and related advantage is that we can “mix and match” the same perturbation and cell representation networks in different ways; for example, after training a network that effectively represents cellular gene expression states, we can combine this same network with a representation network for either small molecules or genetic perturbations without having to re-train the cell representation network. An additional advantage is that the representation networks can be pre-trained on unpaired perturbation and cell observations, which are usually available in much larger quantities than the paired perturbation response data. If a high-quality pre-trained model already exists for encoding a particular type of data, we can directly plug it into the PerturbNet without any further training. The cINN architecture used for the mapping network confers several advantages, including stable and efficient training and a mapping that is invertible by construction (see Methods for details). Additionally, the mapping network can model additional covariates when available, such as dose or cell type, by training the mapping function using both the perturbation representation and the covariates (see Supplementary section A.1).

### PerturbNet predicts response to unseen small molecule treatments

We first investigated whether PerturbNet can predict response to unseen drug treatments. As discussed above, the PerturbNet framework is fully modular, allowing arbitrary network architectures for the representation networks. Thus, we adopted neural network architectures appropriate to the types of data in this prediction task. Because the pharmacological properties of a small molecule are largely determined by its chemical structure, we used a perturbation representation network that can encode molecular structures into low-dimensional vectors. We chose to start from molecules in simplified molecular-input line-entry system (SMILES) format, which describes molecular structures as character strings. We used a previously published architecture (Fig. 2a). We pre-trained the ChemicalVAE on the ZINC dataset [22], which contains 250,000 molecules.

**Fig. 2.**
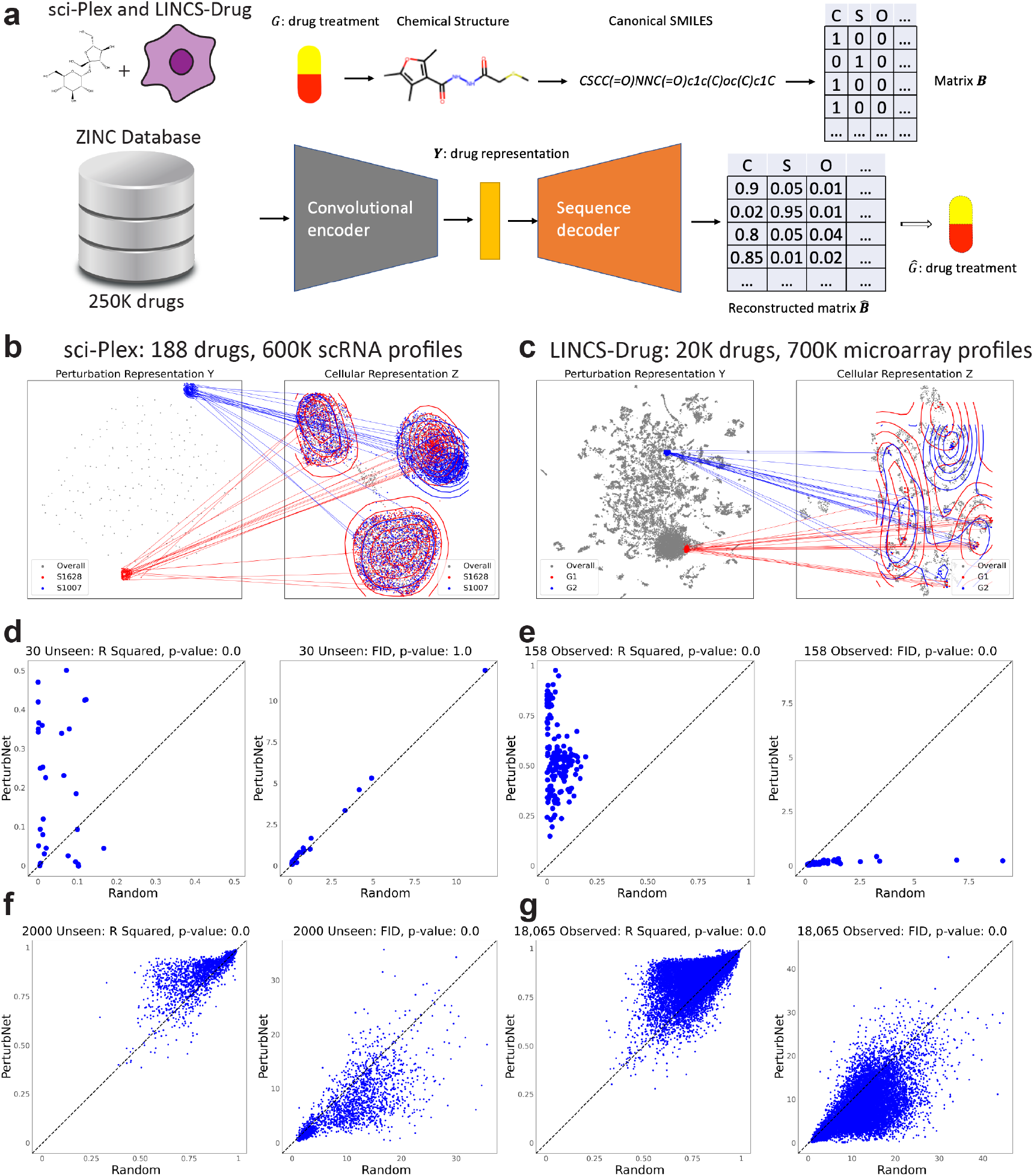
PerturbNet predicts response to small molecule treatment. **a** Diagram of the chemical variational autoencoder (ChemicalVAE) architecture for encoding small molecules represented as SMILES strings. The network was trained on the ZINC dataset, consisting of approximately 250,000 drug-like molecules. **b-c** Visualization of PerturbNet predictions for two distinct perturbations from sci-Plex **b** and LINCS-Drug **c** datasets. The UMAP coordinates are computed from the latent spaces of the perturbation network (left) and cell state network (right). The mapping function learned by the cINN is indicated with lines connecting the perturbation and cell state representations. The predicted cell state distributions are also indicated with contour lines. **d** Scatter plots of R squared and FID metrics for PerturbNet (y-axis) and random baseline (x-axis) for the 30 unseen drug treatments of the sci-Plex dataset. Each point is one held-out drug. The p-values for one-sided Wilcoxon rank-sum tests are shown above each plot. **e** Scatter plots of R squared and FID metrics for the 158 observed drug treatments of the sci-Plex dataset. Each point is one observed drug. **f** Scatter plots of R squared and FID metrics of PerturbNet for the 2000 unseen drug treatments of the LINCS dataset. **g** Scatter plots of R squared and FID metrics for the 18,065 observed drug treatments of the LINCS dataset.

We then trained cell representation networks on two large expression datasets: the Library of Interconnected Network Cell Signatures (LINCS) [23] and sci-Plex [2]. The LINCS dataset consists of 689,831 microarray measurements from 170 different cell lines treated with 20,065 compounds. The sci-Plex dataset contains 648,857 scRNA-seq profiles from three cell lines treated with 188 compounds. We used different types of representation networks for these two types of data: a variational autoencoder (VAE) with Gaussian likelihood for the normalized microarray data and a VAE with negative binomial likelihood for scRNA-seq count data. We used fully-connected layers (multilayer perceptron architectures) for both VAEs. We then trained a cINN for both the LINCS and sci-Plex datasets to translate from the ChemicalVAE latent space to the cell state space. Note that, due to the modular nature of PerturbNet, we were able to use the same ChemicalVAE for both datasets without retraining it.

To visualize the PerturbNet predictions, we plotted UMAP coordinates of the perturbation and cell state spaces. Then we plotted the mapping between the spaces by drawing lines to connect each perturbation with the cell states induced by it and summarizing the cell state distribution using a contour plot. Note that the perturbation representations are probabilistic, so that multiple nearby points on the UMAP plot indicate the distribution of latent representations for each perturbation. These visualizations qualitatively confirm the ability of PerturbNet to model complex perturbation effects, including very different cell state distributions induced by perturbations with distinct representations (Fig. 2b).

To quantitatively evaluate the performance of PerturbNet in predicting effects of unseen small molecule treatments, we held out a subset of the treatments during training. After training, we compared the true and predicted gene expression values for these treatments. We also compared our predictions with a baseline model, in which we randomly sampled cells from the treatments seen during training. Comparison with this baseline is important, because perturbations with very small or no effect can be in principle be accurately predicted simply by guessing the mean of all cells.

We employed the R squared and Fréchet inception distance (FID) metrics to evaluate prediction performance of single-cell responses. The R-squared indicates the slope of the best-fit line between the true and predicted gene expression values for a particular perturbation, and is a measure of how well the mean of the distribution is estimated. The FID is the Wasserstein-2 distance between the true and predicted gene expression distributions, and summarizes the concordance of the expression distributions as a whole. Higher R-squared and lower FID values are better. See Methods for more details.

Overall, PerturbNet predicts the cell states induced by small molecule treatment significantly more accurately than the baseline model (Fig. 2d-g). PerturbNet outperforms the random model at predicting the effects of unseen perturbations (Fig. 2d), achieving significantly better R squared across the 30 held-out perturbations (*p* < 2 × 10^-16^, one-sided Wilcoxon rank-sum test). PerturbNet also significantly outperforms the random model at predicting the 158 observed perturbations in terms of both R squared and FID (Fig. 2e). For the LINCS dataset (Fig. 2f-g), PerturbNet outperforms the random model at predicting the effects of 2000 unseen and 18,065 observed perturbations, achieving significantly better R squared (*p* < 2 × 10^-16^, one-sided Wilcoxon rank-sum test) and FID metrics (*p* < 2 × 10^-16^, one-sided Wilcoxon rank-sum test). The R squared values achieved by PerturbNet are generally higher for the LINCS dataset than the sci-Plex dataset, perhaps because of the significantly larger number of small molecules available for training the cINN (18,065 for LINCS vs. 158 for sci-Plex).

We found that, when covariates are provided, incorporating these can improve PerturbNet performance (Supplementary Section A.1 and Fig. 1). We further explored whether incorporating cellular response data into the perturbation representation network could improve prediction performance (see Supplementary section A.2). To do this, we used the measured gene expression profiles for each perturbation to “fine-tune” the representation network. We calculated the distances between cell state distributions observed for all pairs of perturbations in the training data. Then, we converted these distances to a similarity graph and added a graph regularization term to the VAE loss function. Intuitively, this updated loss function encourages the perturbation representations to simultaneously reconstruct the perturbation from the latent space and preserve the similarity relationship among the perturbations’ cell profiles. Training the cINN with the fine-tuned representation network gave a small but statistically significant improvement in PerturbNet performance for the LINCS-Drug dataset (Supplementary Fig. 3). Interestingly, the KNN model did not show significant improvement when using the fine-tuned latent space.

### PerturbNet predicts response to unseen genetic perturbations

We next extended the PerturbNet framework to genetic perturbations by constructing an autoencoder for the target genes in combinatorial genetic perturbations. Genome editing with CRISPR/Cas9 directly modifies the DNA sequence, leading to changes in the protein-coding sequence or non-coding regulatory sequence. In contrast, genetic perturbations using CRISPR activation (CRISPRa) or CRISPR interference (CRISPRi) do not change the original DNA sequence, but rather change gene expression [10]. These types of perturbations can be described by the identities of their target genes [4]. To encode such genetic perturbations, we developed a new autoencoder that we call GenotypeVAE (Figure 3a). Our key insight is that the numerous functional annotations of each gene (organized into a hierarchy in the gene ontology) provide features for learning a low-dimensional representation of both individual genes and groups of genes. The gene ontology consortium has annotated 18,832 human genes with a total of 15,988 terms (after filtering to remove terms with very low frequency). Using these annotations, we can describe each target gene *g* as a one-hot vector of length 15,988, where a value of 1 in the vector element corresponding to a particular term indicates that the gene has that annotation. If we have a genetic perturbation with multiple target genes, we can simply take the union of the GO annotations from all of the perturbed genes.

**Fig. 3.**
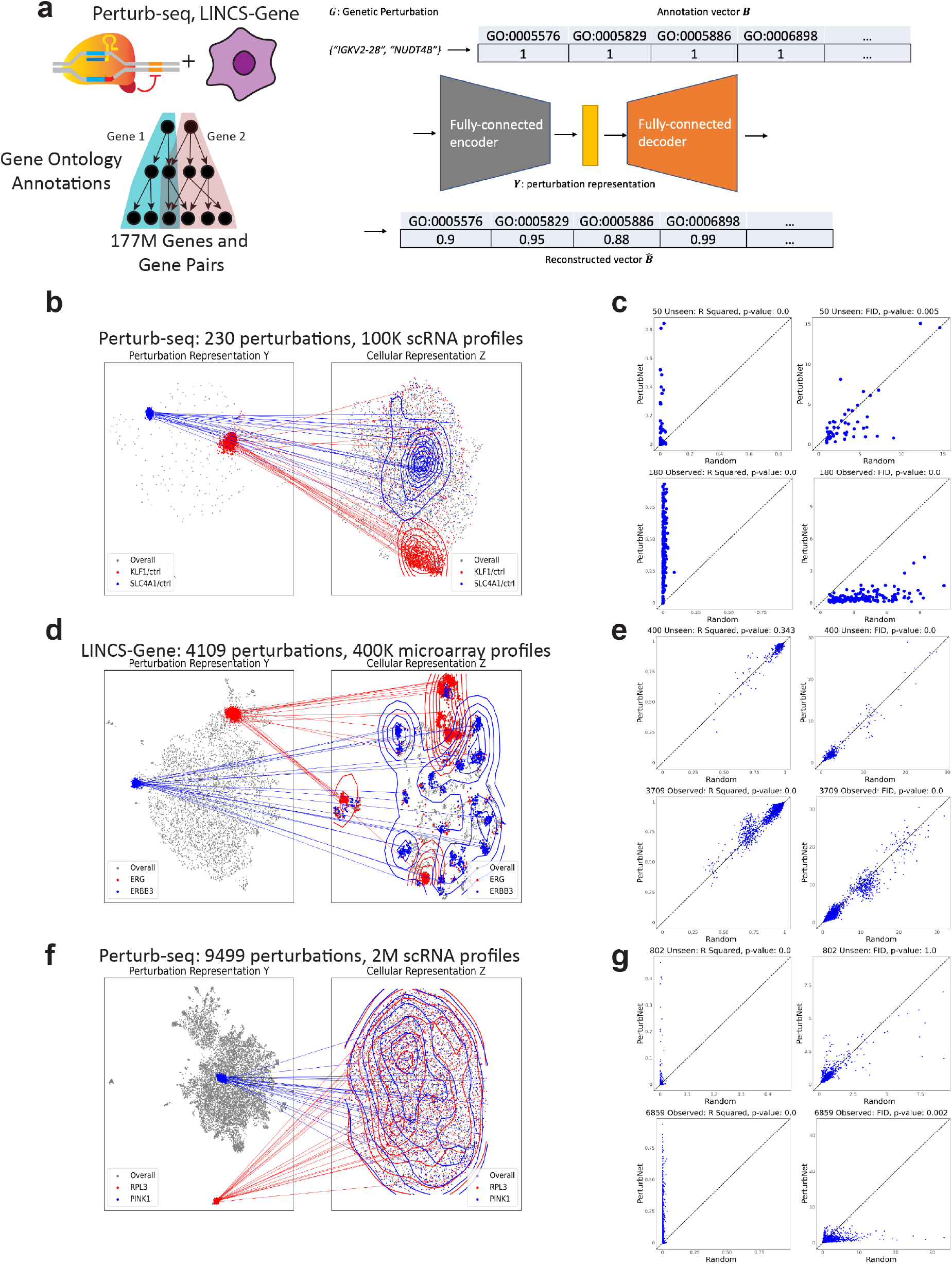
PerturbNet predicts response to genetic perturbation. **a** Diagram of the Geno-typeVAE architecture. Perturbations are represented in terms of their gene ontology annotations. The network is trained on all one- and two-gene combinations (approximately 177 million). **b, d, f** Visualization of PerturbNet predictions for two distinct perturbations from GI **(b)**, LINCS-Gene **(d)**, and GSPS **(f)** datasets. **c, e, g** Scatter plots of R squared and FID metrics for PerturbNet and random baseline for unseen and observed genetic perturbations from the GI **(c)**, LINCS-Gene **(e)**, and GSPS **(g)** datasets. The p-values for one-sided Wilcoxon rank-sum tests are shown above each plot.

We trained a variational autoencoder with fully-connected layers to reconstruct these binary annotation vectors from a latent representation. Our approach is inspired by a previous study that used neural networks to embed genes into a latent space using their gene ontology annotations [24]. We trained GenotypeVAE using one-hot representations of many possible genetic perturbations. Considering all single- and double-gene combinations of the 18,832 human genes with GO term annotations, there are approximately 177 million possible training data points (Fig. 3a).

We evaluated our approach on several large-scale genetic perturbation datasets (Fig. 3b-g). We used data from a CRISPRa screen in K562 cells with 230 perturbations (GI) [10], the LINCS dataset with 4,109 shRNA perturbations (LINCS-Gene) [23], and the genome-scale Perturb-seq dataset with 9499 CRISPRa perturbations (GSPS) [25]. Both GI and GSPS consist of integer count scRNA-seq measurements, while LINCS-Gene consists of real-valued microarray data. We therefore obtained cell state representations by training a VAE with negative binomial likelihood [26] on the GI and GSPS datasets, and a VAE with Gaussian likelihood on the LINCS-Gene dataset.

Visualizing the perturbation representations, cell representations, and mapping functions shows that the PerturbNet can model the distinct cell state distributions induced by different perturbations (Fig. 3b, d, f). For example, two perturbations in the GI dataset (KLF1/ctrl, SLC4A1/ctrl) in the GI dataset (Fig. 3b) and the two knockdowns (ERG, ERBB3) in the LINCS-Gene dataset (Fig. 3d) have very different perturbation representations and cell state distributions. The perturbation representations of the pair of (RPL3, PINK1) in the GSPS data show distinctive distributions (Fig. 3f), while the difference between their cellular representations is less obvious, possibly due in part to batch effects [25].

We predicted single-cell responses to each genetic perturbation using the baseline models and PerturbNet for the three datasets. Fig. 3c show the performance of predicted cell samples evaluated with R squared and FID metrics of PerturbNet over random on the GI data. PerturbNet significantly outperforms the random model for the 180 observed perturbations, and is also significantly better than random for unseen perturbations in R squared.

Fig. 3e shows the prediction performance of PerturbNet compared to the random baseline for the LINCS-Gene data. PerturbNet has significantly lower FID than the random model for both unseen and observed perturbations, and also has higher R squared for the observed perturbations. The random model shows very high R squared values (around 0.75) for the LINCS-Gene data, possibly because most genetic perturbations in the LINCS-Gene dataset have small perturbation effects.

We evaluated the predictive models on the GSPS data [25], with a large number of target genes and a substantial proportion of perturbations with very few cells. We filtered out the genetic perturbations with fewer than or equal to 100 cells before evaluating the baseline models. Fig. 3g shows the performance of the models on the 802 unseen and 6859 observed genetic perturbations with more than 100 cells. PerturbNet has significantly higher R squared than random for both unseen and observed perturbations. However, it does not show better FID than random, possibly due to complex batch effects, as noted by the authors [25].

We also compared the performance between KNN and PerturbNet for the GI, LINCS-Gene and GSPS datasets (Supplementary Fig. 4). PerturbNet shows better predictions than KNN for the observed perturbations of the three datasets. PerturbNet also gives significantly better R squared and FID for unseen perturbations of the LINCS-Gene data, and better FID for unseen perturbations of the GI and GSPS data.

We also fine-tuned GenotypeVAE using the LINCS-Gene data following similar steps as described for the ChemicalVAE fine-tuning above (see Supplementary section A.2). Fine-tuning GenotypeVAE again results in a small but statistically significant improvement in the performance of PerturbNet (Fig. 4). As with the ChemicalVAE, the fine-tuning algorithm improves only the PerturbNet, but does not significantly improve the performance of the KNN model.

**Fig. 4.**
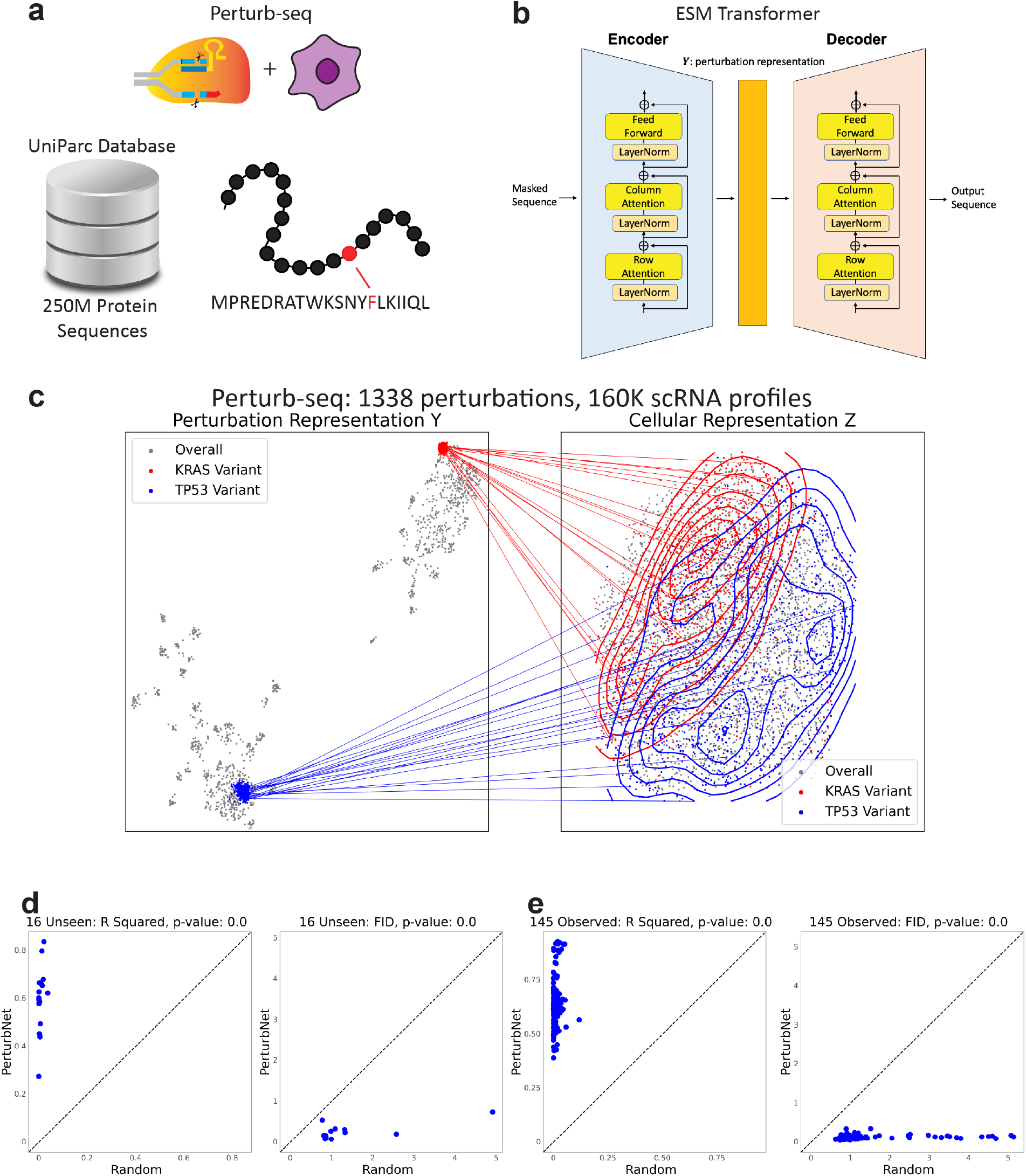
PerturbNet predicts response to coding sequence mutation. **a** Diagram of the approach for training representation network for coding sequence mutations. Each perturbation is an amino acid sequence edited by CRISPR. We used a model pre-trained on the UniParc database, containing 250M sequences. **b** Architecture of the evolutionary scale modeling (ESM) transformer used to embed protein sequences. **c** Visualization of PerturbNet predictions for two distinct perturbations from the Ursu dataset. The UMAP coordinates are computed from the latent spaces of the perturbation network (left) and cell state network (right). The mapping function learned by the cINN is indicated with lines connecting the perturbation and cell state representations. The predicted cell state distributions are also indicated with contour lines. **d, e** Scatter plots of R squared and FID metrics for PerturbNet and random baseline for unseen and observed genetic perturbations from the Ursu dataset. The p-values for one-sided Wilcoxon rank-sum tests are shown above each plot.

### PerturbNet predicts response to coding sequence mutation

In addition to CRISPRi and CRISPRa, which do not change the DNA sequence within a cell, Perturb-seq can also be combined with CRISPR genome editing. For example, Ursu et al. recently used a multiplex CRISPR screen to introduce many different coding sequence mutations into the TP53 and KRAS genes of A549 cells [8]. This study found that the genome edits caused “a functional gradient of states” with continuously varying gene expression profiles. Unlike CRISPRa or CRISPRi perturbations, which can be represented in terms of the identities of the target genes, genome editing perturbations are best represented as the distinct amino acid sequences of either the wild-type or edited genes.

To extend PerturbNet for predicting single-cell gene expression responses to coding sequence variants, we developed a strategy for embedding amino acid sequences. We chose to use the pretrained evolutionary scale modeling (ESM) network [27] to obtain latent representations of the unique protein sequences produced by genome editing (Fig. 4a-b). ESM is a previously published network that was pre-trained on about 250 million protein sequences from the UniParc database [27]. Unlike the chemicalVAE and the GenotypeVAE used above, the ESM encodings are deterministic for a given input sequence; thus, to avoid overfitting when training the cINN on ESM representations, we added a small amount of Gaussian noise sampled from 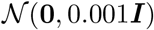.

We then trained PerturbNet on the Ursu dataset, which measured the effects of many distinct mutations in the TP53 and KRAS genes [8]. We preprocessed the detected CRISPR guide RNA sequences to obtain a single, complete protein sequence label for each individual cell. This gave a total of 1,338 unique protein coding sequences observed in the Perturb-Seq data; the vast majority of these unique combinations occur only in one or a handful of cells. We trained a variational autoencoder with negative binomial likelihood on the whole Ursu dataset to obtain latent representations of cell state. Then we trained a cINN to map from protein sequence representation space to cell state space, holding out 130 perturbations. Visualizing the perturbation representations, cellular representations, and mapping function shows that PerturbNet can model the distinct cell state distributions induced by different sequence mutations (Fig. 4c).

Evaluating PerturbNet predictions on the observed and held-out perturbations shows that the model performs significantly better than the random baseline in terms of both R squared and FID metrics (Fig. 4d-e). Note that we filtered the variants to those with more than 400 cells when performing this comparison, because most of the unique protein sequences occur in only one or a few cells. PerturbNet also shows better predictions than KNN for the observed perturbations, and has better FID than KNN for the unseen perturbations (Supplementary Fig. 5d).

### Attributing Perturbation Effects to Specific Perturbation Features

Having established that PerturbNet can successfully predict the effects of unseen perturbations, we reasoned that the model could give insights into which specific perturbation features are most predictive of cell state distribution shifts. For example, it would be desirable to know which atoms in a drug or which gene functions most strongly influence the model predictions. Such insights can give hints about mechanisms and help build confidence that the model is learning meaningful relationships between perturbations and cell states.

We employ the method of integrated gradients [28] to determine, for each feature of a perturbation, whether the presence of the feature increases or decreases the probability of cells being in a particular state. To do this, we divide the space of observed cell states into discrete cell types through unsupervised clustering. Then we train neural networks to classify cell states (including predicted cell states output by the mapping network) into these discrete types (Fig. 5a). For ease of interpretation, we train a binary classifier for each cell type, to classify the cells as either belonging to that type (1) or not (0). We then use the method of integrated gradients to calculate an attribution score for each feature of an input perturbation. This score tells whether each feature increases or decreases the probability of generating cells of a particular type.

**Fig. 5.**
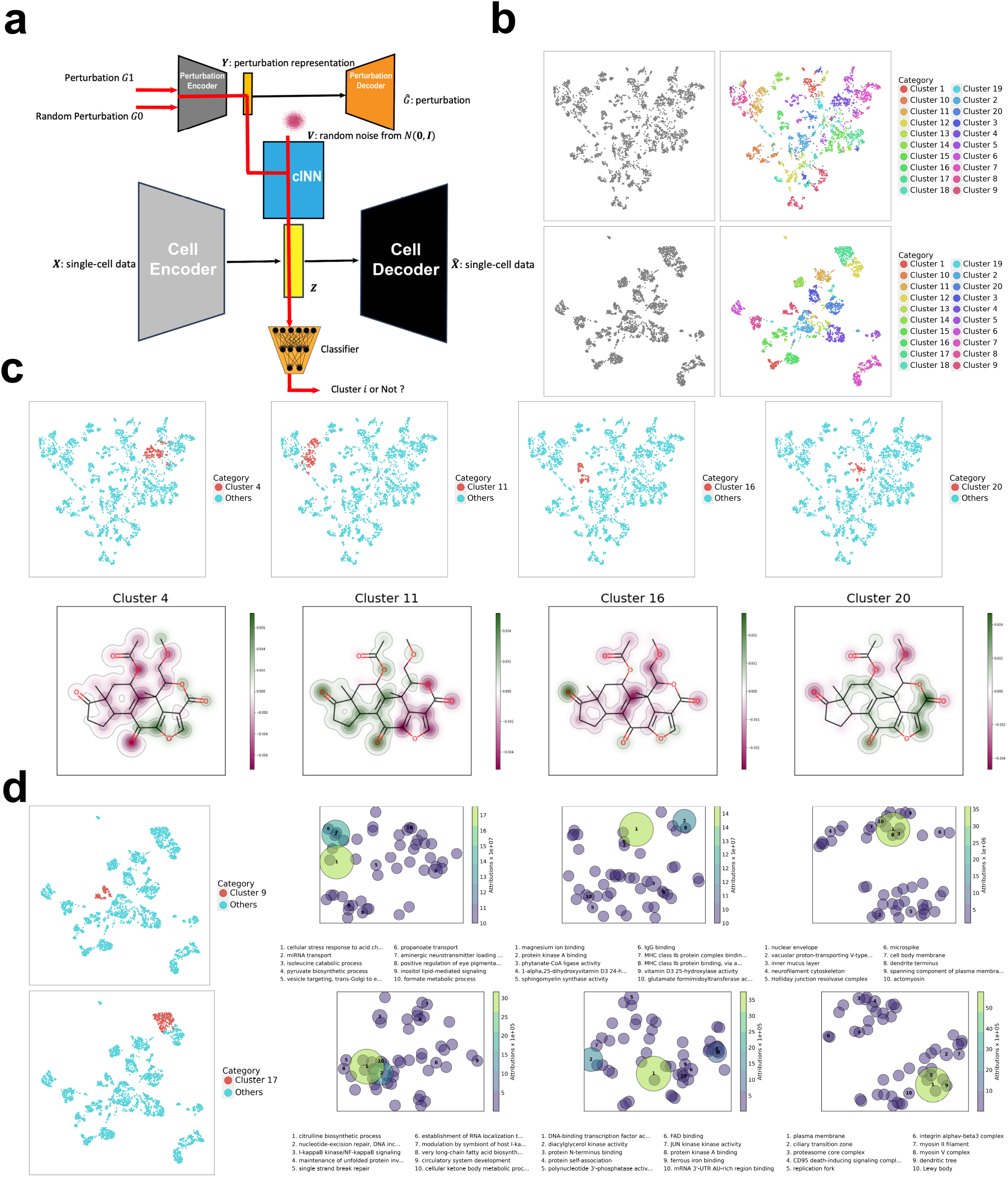
Attributing cell state shifts to specific features of perturbations. **a** Diagram of approach for attributing perturbation outcomes to specific perturbation features. We attach a binary classifier after the cINN to classify cells into discrete types. By comparing the classification results of an input perturbation and a baseline (random perturbations), we can determine which perturbation features increase classification probability. **b** UMAP plots of LINCS-Drug and LINCS-Gene colored by cluster label. **c** UMAP plots of LINCS-Drug with selected clusters and the selected drug colored by attribution scores for each atom. **d** UMAP plots of LINCS-Gene with selected clusters and the GO terms for the selected genetic perturbation. The GO terms are arranged using multidimensional scaling and colored by attribution score.

As an example, we performed integrated gradient attribution on the LINCS-Drug and LINCS-Gene datasets. We performed k-means clustering separately on the latent values of VAEs trained on the LINCS-Drug and LINCS-Gene datasets. In both cases, we divided the cell states for observed cells into *k* = 20 clusters (Figure 5a-b). We then trained neural networks to classify cell latent values into these 20 clusters. For each cluster, we could then calculate an attribution score for each input feature of a perturbation. A positive attribution score indicates that a feature increases the probability of generating cells in that particular cluster, whereas a negative score indicates a decreased probability of generating cells in that cluster.

We visualized the attribution scores for small molecule perturbations by coloring each atom in the molecular structure (Fig. 5c). Results are shown for a representative drug and four different clusters of cell states from the LINCS-Drug dataset. Each cluster has a different attribution pattern; for example, the three-ring structure in the lower-right of the molecule has a positive attribution for clusters 4 and 20 but a negative attribution for cluster 11. Examples of attributions for an additional 12 molecules are shown in Supplementary Fig. 6.

Similarly, we visualized the attribution scores for genetic perturbations (Fig. 5c). For the genetic perturbations, each feature is a gene ontology annotation term. We arrange the terms with top attribution using 2D multidimensional scaling on the GO terms and separately plot the GO terms related to function, process, and component. Cluster 9 attributions implicate cellular stress response and several terms related to protein-ligand binding. Cluster 17 attributions implicate functions related to DNA repair and kinase activity. Examples of attributions for additional clusters are shown in Supplementary Figs. 7-8.

### Designing Perturbations to Achieve Target Cell State Distributions

The ability of PerturbNet to predict out-of-distribution cellular responses can, in principle, be used to design perturbations that achieve a desired outcome. For instance, a search for small molecules that shift cells away from a pathological state could assist with drug discovery. Similarly, predicting genetic perturbations that shift cells toward a target cell state could help improve somatic cell reprogramming protocols. Both applications rely on the notion of counterfactual prediction: predicting what a particular cell would look like if treated with a different perturbation than the one observed.

Fig. 6a gives a high-level summary of the procedure to design perturbations using PerturbNet. We can encode observed cells into the cell state space using the encoder of the cell representation network. Then, using the mapping network in reverse, we can search over the perturbation latent space until we find the perturbation whose cell state distribution most closely matches the target.

**Fig. 6.**
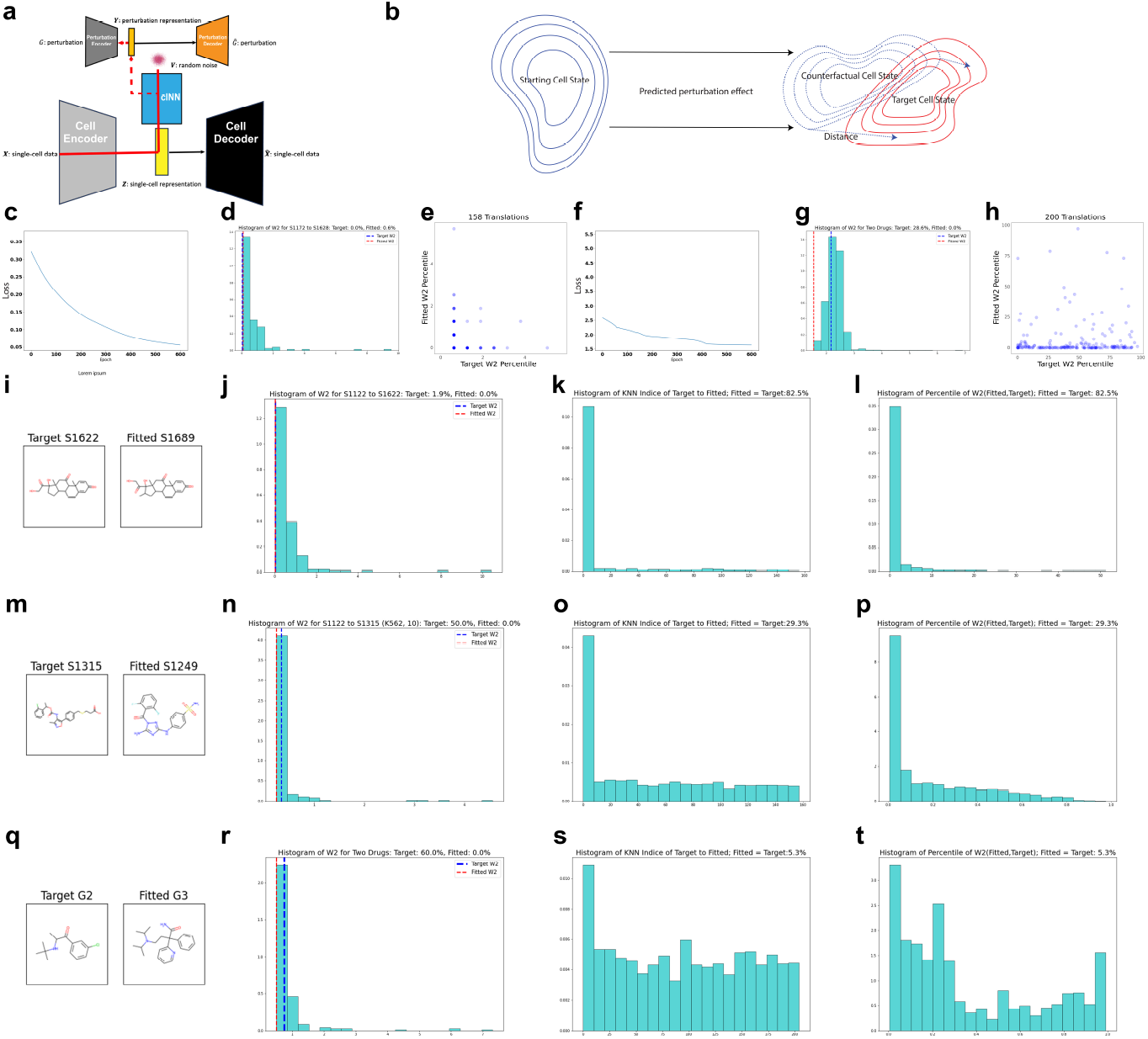
Leveraging PerturbNet to design perturbations with desired outcomes. **a** Diagram of perturbation design strategy. A target cell state distribution is defined by encoding observed cells into the cell state space. The cINN is used in reverse, then optimization is performed in the perturbation space until a perturbation matching the desired distribution is identified. **b** Diagram of the relationship among starting cell state distribution, the counterfactual cell state induced by perturbing the starting cell state, and the target cell state. **c-e** Perturbation design results for sci-Plex using continuous latent space. **c** Example loss curve showing convergence of W2 distance during optimization. **d** Histogram of W2 distance between target cell state and counterfactual (fitted) cell state when using all observed drugs for example perturbation design task. Distances for true drug (target) and fitted drug are indicated as dotted lines. **e** Scatter plot of counterfactual (fitted) vs. target W2 distance percentile (calculated using all observed drugs) for all 158 tested perturbation design tasks (translations). Ideally the fitted percentile should be no larger than the target percentile. **f-h** Same plots as c-e, but for LINCS-Drug. **i-l** Perturbation design results for sci-Plex using latent space locations from observed drugs only. **i** Molecular structures for target and fitted drugs for an example perturbation design task. **j** Histogram of W2 distances as in d, but with fitted perturbation determined from the discrete set of observed perturbations. **k** Histogram of KNN indices across perturbation design tasks. A smaller KNN index means that the fitted perturbation was closer to the true perturbation. The true drug was identified 82.5% of the time. **l** Histogram of W2 distance percentiles across perturbation design tasks. A smaller value means that the fitted perturbation was closer to the true perturbation. The true drug was identified 82.5% of the time. **m-p** Same as i-l but for sci-Plex with covariate adjustment. **q-t** Same as i-l but for LINCS-Drug.

In more detail, the mapping network of PerturbNet allows counterfactual prediction because the mapping function is invertible. Starting from a particular cell state *c*_1_ and perturbation *p*_1_, the cINN can calculate the residual variable *v* that uniquely identifies this combination of state and perturbation. If we then give *v* and a different perturbation *p*_2_ as input to the cINN mapping network, the output cell state *c*_2_ corresponds to the counterfactual state of *c*_1_ under the different perturbation *p*_2_.

We can use this counterfactual prediction capability to identify perturbations that achieve a desired shift in cell state distribution. More formally, consider a starting cell state distribution with latent space values *τ*_1_(***Z***), and a target cell state distribution with latent space values *τ*_2_(***Z***). We want to find a perturbation that changes the cells in the starting cell state to the target cell state. From PerturbNet trained with single-cell perturbation responses, we can obtain the encoded representations for *m* cells in the starting cell state with the latent values {***z***_1_,…, ***z***_*m*_} ~ *τ*_1_(***Z***). Each starting cell is originally treated with a perturbation. For simplicity, we assume that these cells are treated with the same perturbation *p*_1_. The target cell state can be represented by the latent values of *n* cells {***z***_*m*+1_,…, ***z***_*m*+*n*_} ~ *τ*_2_(***Z***). Our goal is to find an alternative perturbation *p*\* that will cause starting cells to shift as close to *τ*_2_(***Z***) as possible.

To find such a perturbation, we first define a measure of the dissimilarity of two cell state distributions. We use Wasserstein-2 (W2) distance, also known as Fréchet distance, to quantify the dissimilarity between the cell state distributions of *τ*_2_(***Z***) and *τ*\*(***Z***). This is a widely used metric for comparing distributions and has even been applied several times in the context of comparing scRNA-seq distributions [29, 30, 31].

The problem of designing a perturbation is then to find the perturbation representation that minimizes the squared W2 distance between counterfactual and target distributions. Because the perturbation representation space is continuous, we can perform stochastic gradient descent to efficiently find the latent space location that minimizes this objective function. Alternatively, if we have a finite, discrete set of candidate perturbations, we can minimize the objective function by exhaustive evaluation–that is, by simply predicting the cell state distribution for every perturbation in the candidate set.

We evaluated our perturbation design strategy using the LINCS-Drug and Sci-Plex datasets. To do this, we selected a target set of observed cells, treated with a particular perturbation *p*_2_, then picked another set of starting cells treated with a different perturbation *p*_1_ and tried to design a perturbation *p*\* to shift the starting cells to the target cell state (Fig. 6c-h). When optimizing over the continuous latent space of small molecules, the W2 distance converged rapidly for both datasets (see example loss curves in Fig. 6c,f). To assess how well the cell state distribution induced by the designed perturbations *p*\* matched the target distribution, we calculated the W2 distances between the target distribution and the distribution induced by *p*\*, as well as every other perturbation in the training dataset. We then calculated the percentile of the W2 distance from *p*\* within the overall distribution of W2 distances from observed perturbations. In most cases, the distribution induced by *p*\* more closely matched the target distribution than most perturbations in the initial dataset, indicating that the optimization procedure is able to effectively identify a latent representation with the desired property (Fig. 6d,e,g,h).

To further evaluate the designed perturbations, we optimized the W2 distance over the discrete space of perturbations for which ground truth cell responses are available (Fig. 6i-t). Constraining the search space in this way allows direct assessment of the true cell responses and molecular structures for the designed perturbations. In particular, if the perturbation design approach is working, we expect the candidate perturbations *p*\* to be similar in structure to the perturbation *p*_2_ that was actually applied to the target cells. For the Sci-Plex (either with and without covariate adjustment) and LINCS-Drug datasets, the optimization procedure produced candidate perturbations *p*\* whose molecular structures were much closer to the perturbation *p*_2_ than expected by chance. In many cases, the designed *p*\* was exactly the same as *p*_2_ (83% for sci-Plex unadjusted, 29% for sci-Plex adjusted, 5% for LINCS-Drug). Even when *p*\* was different from *p*_2_, the molecular structures were often quite similar. For example, although the designed perturbation in Fig. 6i is not identical to the actual perturbation with which the target cells were treated, the two molecules have quite similar structures, with just hydroxyl group and double-bonded oxygen in opposite orientations. Fig 6m shows another example, with two six-atom rings separated by a double bonded oxygen attached to a five-atom ring in both structures. Similarly, the target molecule in Fig. 6q features a six-atom ring adjacent to a double-bonded oxygen atom and an amine group. The chemical structure of *p*\* is more similar to *p*_2_ than expected by chance (Fig. 6k,o,s). In addition, the true cell profiles induced by the candidate perturbations *p*\* were much closer to the target distribution than expected by chance, as measured by W2 distance (Fig. 6l,p,t). In summary, these evaluations suggest that the PerturbNet can be used to design perturbations to approximate a desired cell state distribution.

## Discussion

Our results open a number of exciting future directions. One possible direction is to broaden to additional types of perturbations and responses. For example, one could try to predict the trajectories of single-cell responses after a sequence of perturbations [32]. The PerturbNet might be improved to sequentially model new cellular representation on the new perturbation representation and the previous cellular representation.

Future work could also employ other state-of-the-art methods for chemical and genetic perturbations to obtain better perturbation representations. We can also consider training these frameworks on larger chemical databases such as PubChem [33] or larger GO annotation sets by incorporating genetic perturbations with more than two target genes as well. From our experiments, we find that the prediction performance of PerturbNet on cellular responses to unseen perturbations is likely to be impacted by the number of observed perturbations for training the cINN. To improve the cINN translations for single-cell data with a small number of observed perturbations, one may consider transfer learning [34] to utilize a cINN model trained on a dataset with a large number of perturbations such as LINCS-Drug and LINCS-Gene. In addition, we could incorporate more sophisticated generative models like MichiGAN [35] into the PerturbNet framework. For example, adding an extra training stage to replace the cell state VAE with a conditional generative adversarial network (GAN) could further improve generation performance. We hope that PerturbNet and related approaches can help shape the design of high-throughput perturbation experiments, better leverage these datasets, and ultimately help identify new chemical and genetic therapies.

## Acknowledgements

We thank Hojae Lee and Yuwei Bao for helpful discussions and Joseph Replogle for sharing the genome-scale Perturb-seq data. This work was supported by NIH grants R01HG101883 and U01HG011952 to J.D.W.

## Author contributions

H.Y. and J.D.W. conceived the idea of PerturbNet. H.Y. implemented the approach and performed data analyses. H.Y. and J.D.W. wrote the paper.

## Data Availability

All datasets analyzed here are previously published and freely available.

## Code Availability

PerturbNet code is available on GitHub: https://github.com/welch-lab/PerturbNet

## Materials & Correspondence

Please address correspondence to Joshua D. Welch.

## Methods

### Datasets with chemical perturbations

We have the ZINC data to train the ChemicalVAE model. We utilize the sci-Plex and LINCS-Drug data with cellular responses to chemical perturbations.

**Table 1:** High-Throughput Gene Expression Datasets with Chemical Perturbations.

| Dataset | sci-Plex | LINCS-Drug |
| --- | --- | --- |
| Source | scRNA-seq | Microarrays |
| Cell Lines | A549, K562, MCF7 | ~100 |
| Number of Measurements | 648,857 | 689,831 |
| Number of Genes | 5087 | 978 |
| Number of Perturbations | 188 | 20,065 |

#### ZINC

We obtained the ZINC database with 250,000 compounds [22] from the ChemicalVAE model (https://github.com/aspuru-guzik-group/chemical_vae/tree/main/models/zinc). We transformed the compounds to canonical SMILES following the ChemicalVAE tutorial (https://github.com/aspuru-guzik-group/chemical_vae/blob/main/examples/intro_to_chemvae.ipynb) via the RDKit package [36]. We also utilized the chemical elements’ library from this tutorial to define the one-hot matrices of drug treatments, where we constrained the maximum length of canonical SMILES strings to be 120.

#### sci-Plex

We processed the whole sci-Plex data [2] using SCANPY [37] with a total of 648,857 cells and 5087 genes. There were 634,110 cells perturbed by 188 drug treatments in total, with 14,627 cells with no SMILES string and 120 unperturbed cells. We randomly selected 30 drug treatments as unseen perturbations and the other 158 drug treatments as observed perturbations.

#### LINCS-Drug

We obtained the LINCS dataset [23] from GEO accession ID GSE92742. The LINCS data had been processed with 1,319,138 cells and 978 landmark genes, containing the LINCS-Drug subset with 689,831 cells treated by 20,329 drug treatments denoted with their SMILES, 20,065 drug treatments of which had lengths smaller than 120. We randomly selected 2000 drug treatments as unseen perturbations and the other 18,065 drug treatments as observed perturbations.

We transformed the SMILES strings of drug treatments of the sci-Plex and LINCS-Drug data to their one-hot matrices according to the chemical elements’ library.

### Datasets with gene knockdowns and coding sequence mutations

We have the GO annotation data to train the GenotypeVAE model. We utilize the GI, LINCS-Gene and GSPS data with cellular responses to gene knockdowns. We use the Ursu data with cellular responses to coding sequence mutations.

**Table 2:** High-throughput gene expression datasets with gene knockdowns and coding sequence mutations.

| Dataset | GI | LINCS-Gene | GSPS | Ursu et al. |
| --- | --- | --- | --- | --- |
| Source | scRNA-seq | Microarrays | scRNA-seq | scRNA-seq |
| Cell Lines | K562 | ~100 | K562 | A549 |
| Number of Measurements | 109,738 | 442,684 | 1,989,373 | 164,931 |
| Number of Genes | 2279 | 978 | 2000 | 1629 |
| Number of Perturbations | 230 | 4109 | 9499 | 1338 |
| Perturbation Identity | Gene | Gene | Gene | Sequence |

#### GO annotations

We obtained the GO annotation dataset for human proteins from the GO Consortium at http://geneontology.org/docs/guide-go-evidence-codes. We removed the annotations of three sources without sufficient information: inferred from electronic annotation (IEA), no biological data available (ND) and non-traceable author statement (NAS). The filtered dataset had 15,988 possible annotations for 18,832 genes.

#### GI

We obtained the GI data on GEO accession ID GSE133344 [10]. Each cell was perturbed with 0, 1 or 2 target genes. We processed the GI data using SCANPY [37] with 109,738 cells and 2279 genes. The processed GI data contained 236 unique genetic perturbations for 105 target genes and 11,726 cells were unperturbed. There were 230 out of 236 genetic perturbations that could be mapped to the GO annotation dataset. We randomly selected 50 genetic perturbations as unseen and the other 180 perturbations as observed.

#### LINCS-Gene

We obtained the LINCS dataset [23] from GEO accession ID GSE92742. The LINCS data had been processed with 1,319,138 cells and 978 landmark genes. The LINCS-Gene subset of the LINCS data contained 442,684 cells treated by 4371 genetic perturbations with single target genes. A total of 4109 out of 4371 genetic perturbations could be mapped to the GO annotation dataset, and we randomly selected 400 genetic perturbations as unseen perturbations and the other 3709 as observed perturbations.

#### GSPS

We used SCANPY to preprocess the GSPS data [25] and to select the top 2000 highly-variable genes with respect to the batches of ‘gemgroups’. The GSPS dataset contained 1,989,373 cells treated by 9867 genetic perturbations with single target genes. There were 9499 genetic perturbations that can be mapped to the GO annotation library. We randomly selected 1000 genetic perturbations as unseen perturbations and the other 8499 as observed perturbations. There were 802 unseen and 6859 observed perturbations, each with more than 100 cells.

#### Ursu et al

We obtained the Ursu data from GEO accession ID GSE161824, and filtered the raw data according to the processed datasets and concatenated the two datasets with KRAS variants and TP53 variants, using their common genes. We preprocessed the concatenated data using SCANPY, containing 164,931 cells and 1629 genes. We also collected the variants from the modifications on the original KRAS and TP53 protein sequences. We obtained 596 KRAS sequences and 742 TP53 protein sequences, and randomly selected 60 KRAS and 70 TP53 variants as unseen perturbations. There were 16 unseen and 145 observed variants with more than 400 cells.

### ChemicalVAE

The commonly used one-hot encoding approach can transform drug treatment labels to a vector of 1’s and 0’s, but it needs pre-specifying the total number of possible drug treatments and cannot encode new treatments after the specification. Therefore, we consider flexible representations ***Y*** for drug treatments to predict drug treatment effects on single-cell data for unseen perturbations.

A drug treatment contains abundant information more than just a label such as ‘S1096’. Its pharmacological properties are usually determined by its chemical structure. We thus aim to encode drugs’ chemical structures to dense representations. We consider drug treatments’ simplified molecular-input line-entry system (SMILES) strings, which distinctively represent chemical structures and treatment information. Although SMILES strings can be encoded to numerical representations through molecular Morgan fingerprints [38] or through language models [39, 40], the representations from these methods are deterministic, meaning that the representations remain the same in replicated encoding implementations. Given that a chemical screen experiment usually contains a limited number of distinct drug treatments, the use of stochastic representations of the drug treatments prevents possible model overfitting.

To improve the learning capacity, especially for representations of unseen treatments, we consider using a chemical variational autoencoder (ChemicalVAE) to generate the stochastic sampled representation ***Y*** of each drug’s SMILES string [41, 42]. In essence, the ChemicalVAE first transforms and standardizes SMILES strings to their canonical forms and tokenizes each canonical SMILES to be encoded as a one-hot matrix. For a canonical SMILES string, the *i*th row of its one-hot matrix corresponds to its *i*th place, and has the *j*th column being 1 and all other columns being 0’s, if its *i*th place has the *j*th character in the collected chemical elements’ library. The one-hot matrices of SMILES strings are then fitted into ChemicalVAE which provides representations ***Y*** for SMILES strings of drug treatments *g.*

We followed the ChemicalVAE model utilized in *Gómez-Bombarelli et al.* (2018) [43] and adapted it to PyTorch implementations. The ChemicalVAE model takes each input of size of 120 by 35, and has three one-dimensional convolution layers with the triplet of number of input channels, number of output channels and kernel size being (120, 9, 9), (9, 9, 9) and (9, 10, 11), respectively. There are a Tanh activation function and a batch normalization layer following each convolution layer. After these transformations, the input is then flattened to a fully-connected (FC) hidden layer with 196 neurons, and is subsequently activated by a Tanh function, followed by a dropout regularization with a dropout probability of 0.08 and a batch normalization layer. Then two hidden layers both with 196 neurons generate means and standard deviations of the latent variable. The decoder of the ChemicalVAE model has a FC hidden layer with 196 neurons, followed by a Tanh activation, a dropout regularization with a dropout probability of 0.08 and a batch normalization layer. Then the elements of the input are repeated 120 times to be put in a GRU layer with three hidden layers of 488 hidden neurons, followed by a Tanh activation. The input is then transformed to a two-dimensional tensor to be put in a FC layer with 35 neurons and a softmax activation function. Then each input is reshaped to be the output tensor of 120 by 35.

We implemented the ChemicalVAE training on the ZINC data with different learning rates. We finally had an optimal training with a batch size of 128 and a learning rate of 10^-4^ for 100 epochs.

### GenotypeVAE

For gene knockdowns, most of the existing methods one-hot-encode the target genes across a set of genes [4] or all genes on a coding sequence [44]. However, this strategy cannot generalize to perturbations with an unseen target gene.

To encode genetic perturbations, we propose a more parsimonious framework and refer to it as GenotypeVAE. Our key insight is that the numerous functional annotations of each gene (organized into a hierarchy in the gene ontology) provide features for learning a low-dimensional representation of individual genes and groups of genes. Using gene ontology (GO) terms, we can represent each target gene *g* as a one-hot vector ***B**_g_*, where 1’s in the vector element correspond to a particular term indicating that the gene has the annotation. Our approach is inspired by *Chicco et al.* (2014) [24]. If we have a genetic perturbation with multiple target genes {*g*_1_,…,*g_k_*}, we use annotation-wise union operations to generate a one-hot annotation vector for the genetic perturbation as follows:

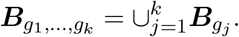

Then, we can train GenotypeVAE using one-hot representations of many possible genetic perturbations. We use the GO Consortium gene ontology annotation dataset of human genes. This resource annotates 18,832 genes with 15,988 annotation terms (after removing some annotations with insufficient information). We take the 15,988-dimensional annotation vector as the input to the GenotypeVAE encoder consisting of two hidden layers with 512 and 256 neurons, following output layers for means and standard deviations, both with 10 neurons. The GenotypeVAE decoder also has two hidden layers with 256 and 512 neurons, along with an output layer of 15,988 neurons activated by the sigmoid activation function. We also have a batch normalization layer, Leaky Rectified Linear Unit (ReLU) activation and a dropout layer with a dropout probability of 0.2 following each hidden layer of GenotypeVAE.

We adjusted different learning rates, batch size and epochs. We finally trained GenotypeVAE on the annotation vectors of single and double target genes from the GO annotation dataset with batch size of 128 for 300 epochs at a learning rate of 10^-4^.

### ESM

A coding variant can be uniquely represented by the protein sequence resulting from the nucleotide alterations induced by CRISPR/Cas9 editing. Similar to chemical perturbations, coding variants can also be summarized as sequences of strings. A key difference is that each character of a protein sequence is a naturally occurring character sequence, whereas a chemical structure is actually a three-dimensional structure (even if it is sometimes represented as a string).

We therefore consider a state-of-the-art language model for protein sequences. Rather than designing our own model and training it from scratch, we employ the previously published Evolutionary Scale Modeling (ESM) [27] architecture. ESM is a self-supervised transformer model [45] and was previously shown to achieve better representations and prediction performance on protein sequences compared to other language models such as long short-term memory (LSTM) networks. As with other transformer models [46], the ESM model was pre-trained on large protein sequence datasets [47]. We adopt a pre-trained ESM model specialized for prediction of single variant effects [48], because this application is most similar to our scenario.

However, the representation obtained from ESM is deterministic for a given protein sequence. The fixed protein representations limit the amount of training data available for PerturbNet, especially when there is a small number of protein sequences. We therefore add low-variance noise ***ϵ*** to the ESM representation ***Y***_ESM_ from ESM. The final perturbation representation is thus computed as

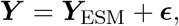

where 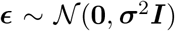. We choose the variance ***σ***^2^ to be a positive constant small enough that it does not significantly alter the relative distances between proteins in the ESM latent space.

### KNN model

From the perturbation representations, ***Y***, of drug treatments, we can learn the relationship of several drug treatments in their latent space. We assume that drug treatments with close latent values tend to also have similar single-cell responses. Thus, the distributions of perturbation responses *p*(***X***|*G* = *g*_1_) and *p*(***X***|*G* = *g*_2_) are similar if *g*_1_ and *g*_2_ have close representations of ***y***_1_ and ***y***_2_.

We then propose our baseline model using the *k*-nearest neighbors (KNN) algorithm to predict single-cell data under drug treatments in Algorithm 1. From ChemicalVAE, we can obtain the representation ***Y*** for a set of treatments 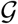, each of which has measured single-cell samples. Then for a drug treatment 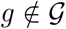 with representation ***y***, we can find its *k* nearest neighbors {*g*_(1)_,…, *g*_(*k*)_} from 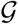 based on ***Y***. We then sample single-cell samples treated with the k nearest treatments in proportion to their exponentiated negative distances to the treatment of interest in the latent space of ***Y***. The sampled single-cell data can be regarded as a baseline prediction for the single-cell data with the treatment of interest.

**Algorithm 1:**
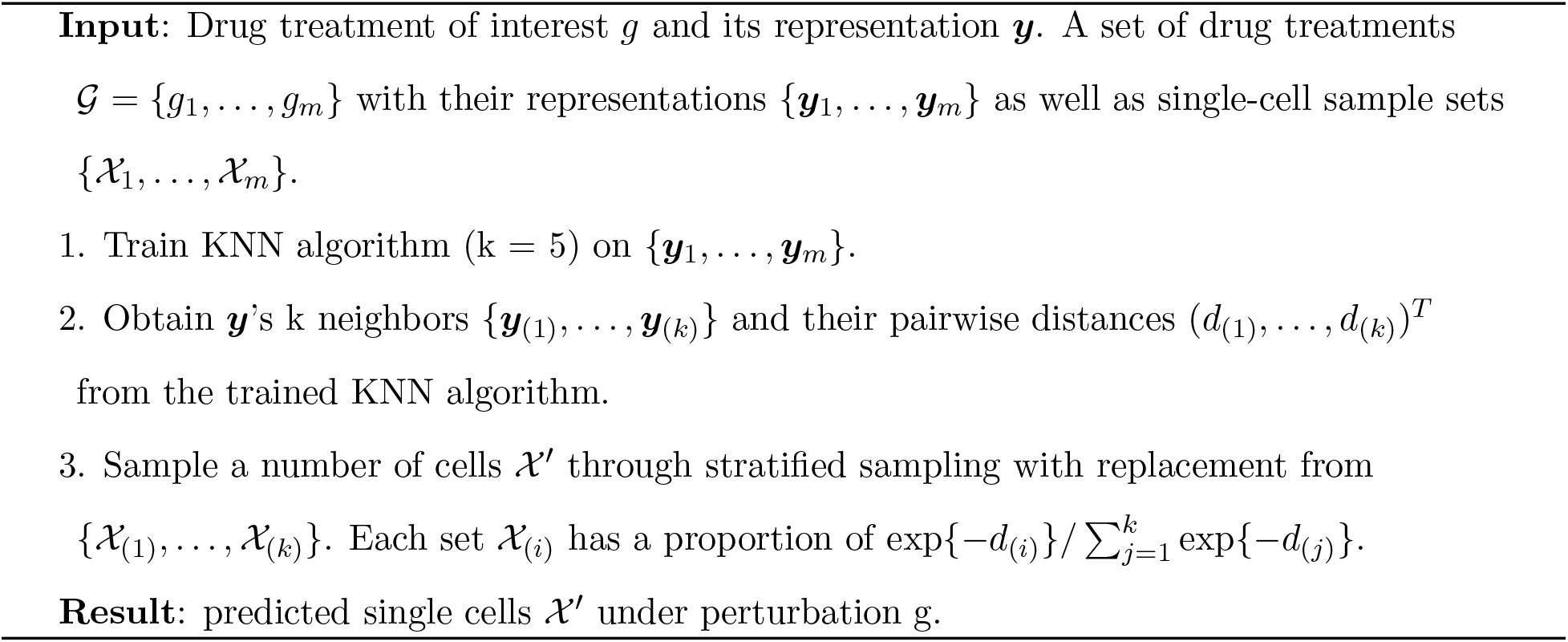
Baseline KNN Model

### Baseline random model

The key assumption of the baseline KNN model is that the perturbation representation is informative to infer single-cell data. To test the informativeness assumption on the perturbation representation, we propose a naive baseline random model in Algorithm 2 that randomly samples single-cell samples under treatments other than the target treatment. If the perturbation representation is uninformative to inferring cell state or cellular response, the random model is likely to have a similar performance to the KNN model.

**Algorithm 2:**
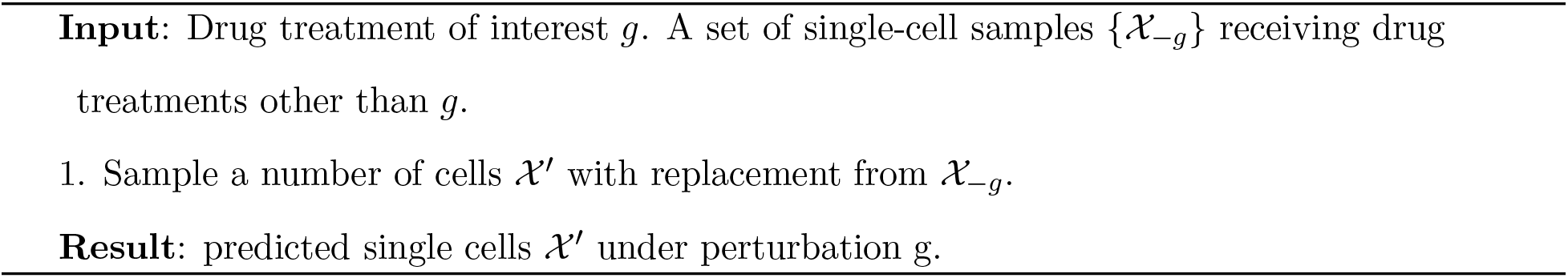
Baseline Random Model

### Prediction metrics

We utilize metrics of R squared and FID score to evaluate the prediction metrics of different models.

#### R squared

We follow the R Squared metric utilized in several frameworks to predict single-cell responses to perturbations [11, 13, 16]. We first obtain the normalized data of predicted and real single-cell responses to a perturbation for the sci-Plex data. We conduct similar processing steps to SCANPY [37]. We first normalize the total number of counts of each cell to be 10^4^, take log-transformation, and scale the values. We directly use LINCS samples as they have already been normalized. We compute the mean gene expression values of normalized data of both predicted and real cells to a drug treatment. We then fit a simple linear regression model on the real mean gene expression values over the predicted mean gene expression values. The R squared of the fitted linear regression is then reported to quantify the accuracy of predicted cells.

#### FID score

We define an FID score metric similar to the FID metric utilized in image data [49]. We train a single-cell VAE model on the whole single-cell dataset using either negative binomial or Gaussian likelihood depending on the data type. We obtain the cell latent values of the predicted and real cells to a perturbation. We then apply the Fréchet distance to the latent values of predicted and real cells with the Gaussian assumption

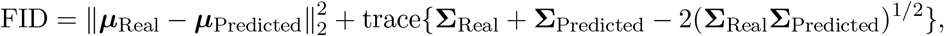

where ***μ***_Real_, ***μ***_Predicted_ are means of predicted and real latent values, and **∑**_Rea1_, **∑**_Predicted_ are co-variance matrices of predicted and real latent values.

### Conditional invertible neural network (cINN)

We consider employing complex normalizing flows of invertible neural networks to understand the relationship between perturbation representation and cellular responses. An affine coupling block [50] enables the input 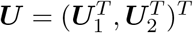 to be transformed to output 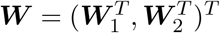 with:

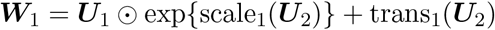

and

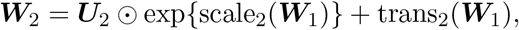

where scale_1_(·), scale_2_(·), trans_1_(·), trans_2_(·) are arbitrary scale and transformation neural networks, and ⊙ is the Hadamard product or element-wise product. The inverse of the coupling blocking can be represented by

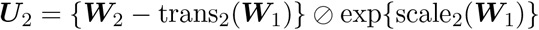

and

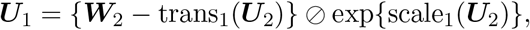

where ⊘ is the element-wise division. The affine coupling block allows bijective transformations between ***U*** and ***W*** with strictly upper or lower triangular Jacobian matrices. A conditional coupling block is further adapted to concatenate a conditioning variable with inputs in scale and transformation networks. A conditional coupling block preserves the invertibility of the block and the simplicity of the Jacobian determinant.

A conditional invertible neural network (cINN) [51, 21] is a type of conditional normalizing flow with conditional coupling blocks and activation normalization (actnorm) layers [52], with both forward and inverse translations. Denote representations from two domains as 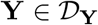 and 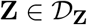. A cINN modeling ***Z*** over ***Y*** gives forward translation

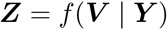

and inverse translation

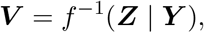

where 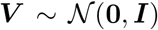. The cINN effectively models *p*(***Z***|***Y***), the probabilistic dependency of ***Z*** over ***Y*** with a residual variable ***V***. As a cINN seeks to extract the shared information from ***Y*** and add residual information ***V*** to generate ***Z***, the objective function to train a cINN is the Kullback-Leibler (KL) divergence between the residual’s posterior *q*(***V***|***Y***) and its prior *p*(***V***). The objective function can further be derived to

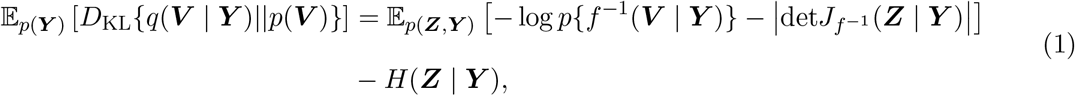

where det J_*f*^-1^_ is the determinant of the Jacobian matrix of *f*^-1^ and *H* is a constant entropy. The optimal *f* that minimizes the objective function in Equation (1) gives *q*(***V***|***Y***) = *p*(***V***). In addition, the objective is an upper bound of the mutual information *I*(***V, Y***). Therefore, a well-trained cINN effectively achieves independence between ***V*** and ***Y***. cINN has the same parameters for forward and inverse translations, reducing the number of model parameters while still preserving network details in both translation directions, and has been utilized to translate domain representations of images and texts [21].

We trained the cINN translations following *Rombach et al.* (2020) [21], where a cINN consists of 20 invertible neural network blocks and an embedding module. Each block has an alternating affine coupling layer, an actnorm layer and a fixed permutation layer. The embedding module consists of FC hidden layers and Leaky ReLU activation functions to embed the conditioning variable into a 10-dimensional variable. We fixed the batch size of 128, the learning rate of 4.5 × 10^-6^ and varied different numbers of epochs for training cINN. We found the cINN training generally stabilized after 50 epochs across different datasets.

### Fine-tuning ChemicalVAE and GenotypeVAE

As both KNN and PerturbNet methods predict cell state based on perturbation representation, it might enhance the prediction performance for cell state from perturbation to use perturbation representation that learns cellular representation information.

We propose an algorithm to fine-tune ChemicalVAE and GenotypeVAE, by adding their evidence lower bound (ELBO) loss with an extra term for a certain cellular property quantity [43]. In this study, we compute the Wasserstein-2 (W2) distance between cellular representations of each pair of perturbations and penalize the trace of ***Y***’s second moment weighted by the Laplacian matrix ***L*** of the adjacency matrix defined from pairwise distances [53]. Denote ***y*** and ***L*** as the perturbation representations and the Laplacian matrix of the perturbations. The regularization term is defined as trace(***y***^*T*^***Ly***), similar to a term commonly arising in spectral graph theory. By penalizing the proposed quantity, we expect perturbations with similar cell states to have closer perturbation representations from ChemicalVAE or GenotypeVAE. To implement the fine-tuning algorithm, we alternate the VAEs’ training with a batch of chemical SMILES strings from a large chemical database or a batch of target genes from the GO annotation dataset with the ELBO loss and another batch of pairs of perturbations and cellular representations from a single-cell chemical or genetic screen dataset with the penalized ELBO loss. We tune a hyperparameter λ on the extra term to adjust the fine-tuning performance. We summarize the ChemicalVAE and GenotypeVAE fine-tuning in Algorithm 3.

**Algorithm 3:**
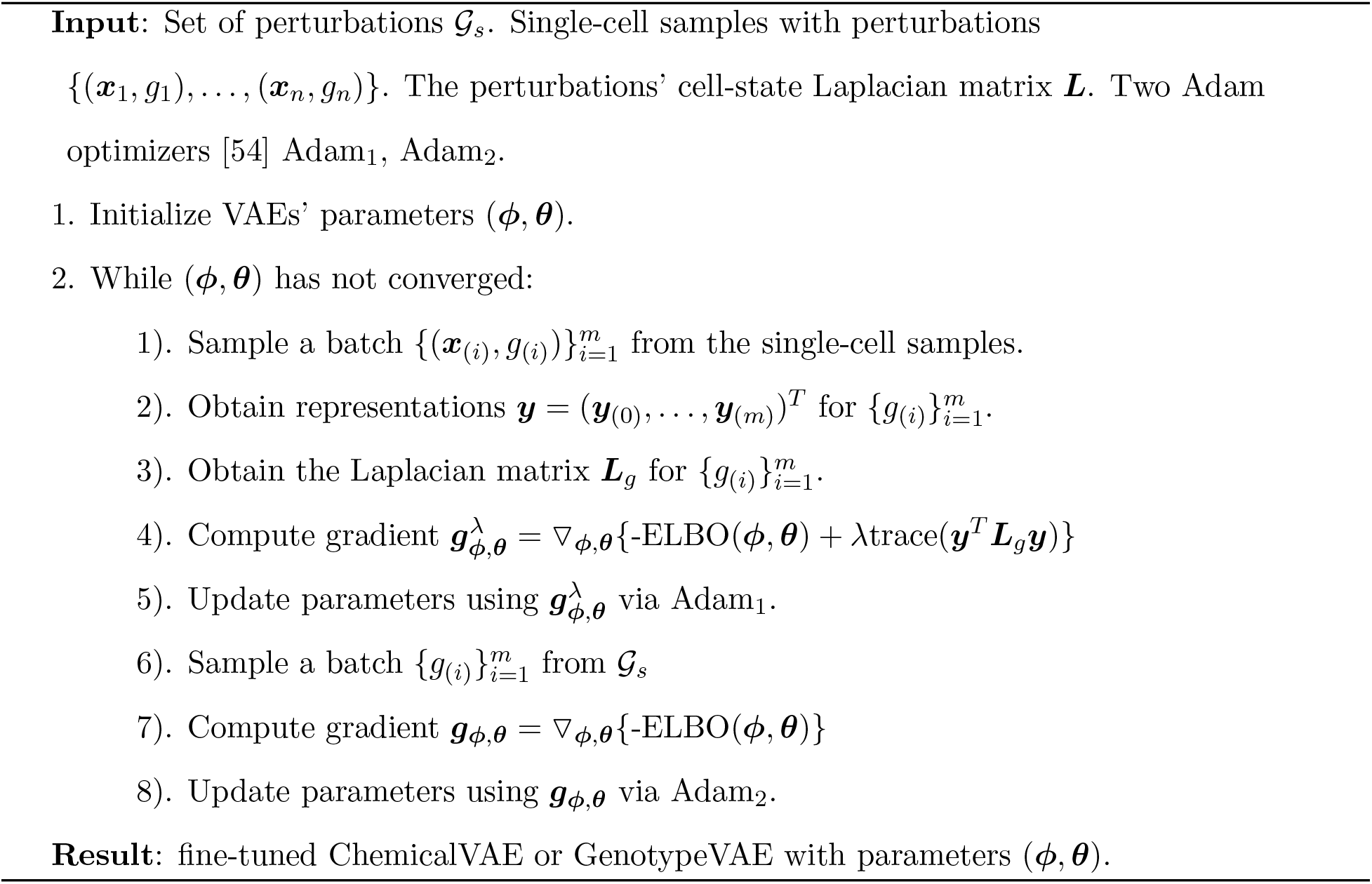
ChemicalVAE and GenotypeVAE fine-tuning

### Optimal perturbation design with continuous and discrete optimizations

Consider a starting cell state with latent space values *τ*_1_(***Z***), and a target cell state with latent space values *τ*_2_(***Z***). We want to find a perturbation that changes the cells in the starting cell state to the target cell state. From PerturbNet trained with single-cell perturbation responses, we can obtain the encoded representations for *m* cells in the starting cell state with the latent values {*z*_1_,…, ***z***_*m*_} ~ *τ*_1_(***Z***). Each starting cell is originally treated with a perturbation. For simplicity, we assume that these cells are treated with the same perturbation *g*_1_. The target cell state can be represented by the latent values of *n* cells {***z***_*m*+1_,…, ***z***_*m*+*n*_} ~ *τ*_2_(***Z***). The optimal translation task thus aims to find an alternative perturbation *g*\* for the starting cells to change their cell state to be close to *τ*_2_(***Z***).

As PerturbNet translates perturbation representation ***Y*** and residual representation ***V*** to cellular representation ***Z***, we can predict the counterfactual cell state under a new perturbation for each cell with two translation procedures. Denote cINN forward translation as *f*(·), ***B***_1_ as the perturbation matrix of the starting perturbation *g*_1_ and the perturbation encoder as *h*(·). First, we obtain residual values {***v***_1_,…, ***v**_m_*} with the inverse translation function *v_i_* = *f*^-1^(***z***_*i*_|***y**_i_*) with perturbation representation ***y**_i_* = *h*(***B***_1_). The translation function then gives each cell’s counter-factual cellular representation ***z**_i_*,* = *f* (***v**_i_*| ***y***\*) under an alternative perturbation’s representation value ***y***\*. We therefore seek the translated counterfactual cell state {***z***_1,*_,…,***z**_m,*_*} ~ *τ*_*_(***Z***) to have a similar distribution to {***z***_*m*+1_,…, ***z***_*m*+*n*_} ~ *τ*_2_(***Z***).

We devise a method to design a perturbation representation ***y***\* that shifts the cells in the starting cell state to approximate the target cell state. To quantify the difference between the cell state distributions, we use Wasserstein distance, which has been widely used to quantify cell populations’ distance [29, 30, 31]. We use Wasserstein-2 (W2) distance [55], which is also known as Fréchet distance, to quantify the dissimilarity between the cell state distributions of *τ*_2_(***Z***) and *τ*_*_(***Z***). The W2 distance is defined as

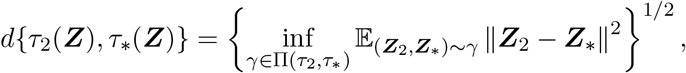

where Π(*τ*_2_, *τ*_*_) is the set of all joint distributions *γ* (***Z***_2_, ***Z***\*) whose marginal distributions are *τ*_2_(***Z***) and *τ*_*_(***Z***), respectively.

Evaluating the W2 distance is extremely difficult for general distributions. To simplify the calculations of the W2 distance, we assume that latent spaces follow multivariate Gaussian distributions [56, 49], which is also commonly assumed in calculating Fréchet inception distance (FID) in image data [49]. Assuming that the latent space *τ_i_*(***Z***) has a multivariate Gaussian distribution 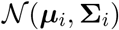 for *i* ∈ {2, *}, the squared W2 distance has a closed form:

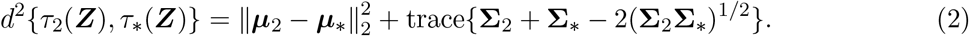

Therefore, we can evaluate the squared W2 distance between the translated counterfactual cell state and the target cell state as 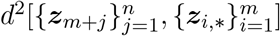. The problem of designing a desired perturbation is then to find the optimal 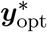 that minimizes the squared W2 distance:

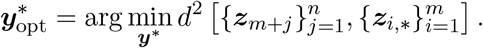

We can further infer the optimal perturbation from representation 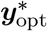. Figure 6a summarizes the procedure to design the optimal perturbation using PerturbNet.

Based on the objective above, we propose what we refer to as “continuous optimal translation.” We first initialize a value for 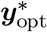 from the standard multivariate Gaussian distribution and then we perform stochastic gradient descent with momentum [54] to minimize the squared W2 loss over 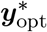. One important implementation detail concerns the calculation of the W2 distance. The distance formula includes the term (**Σ**_2_**Σ**\*)^1/2^, which is difficult to calculate and can become ill-conditioned or approximately singular. We thus rewrite the term as

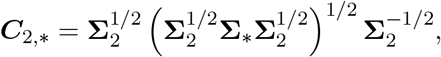

which allows us to replace the difficult term with 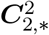 as 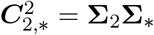.

We use the Adam optimizer to perform stochastic gradient descent with momentum. For the matrix square root terms in ***C***_2,*_, 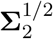 keeps a fixed value during training, and 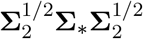 is much more likely than the original term **Σ**_2_**Σ**_*_ to be symmetric positive semi-definite and have a square root matrix. After the continuous optimization, we obtain an optimal perturbation representation 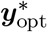 that represents a potential perturbation that achieves the desired shift in cell state distribution.

The continuous optimal translation model can give an optimal perturbation representation 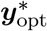 that translates the starting cells to have a similar cell state to *τ*_2_(***Z***). If the real cells in the target cell state {***z***_*m*+1_,…, ***z***_*m*+*n*_} ~ *τ*_2_(***Z***) are treated by a perturbation *g*_2_, we can compare it with the fitted optimal perturbation representation 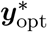 to evaluate if the optimal perturbation representation can achieve the desired cell state shift like the perturbation *g*_2_.

However, the chemical or genetic perturbation from the optimal perturbation representation of a continuous optimal translation is not immediately clear, as an inference model needs to be processed on the perturbation representation. Although it is possible to employ the perturbation generative model to generate chemical or genetic perturbations, doing so brings a host of additional challenges related to molecular structure optimization [43], which is not the focus of this study.

To design the optimal perturbation to achieve the desired cell state shift, we propose another perturbation design strategy that uses discrete optimization. Rather than optimizing the squared W2 loss in the continuous space, the discrete optimal translation searches through a constrained set 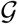 of perturbations, and calculates the squared W2 distance 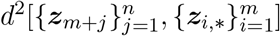 for each perturbation 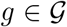 with ***y***\* = *h*(***B**_g_*). Then the optimal perturbation is selected as the one giving the smallest distance so that

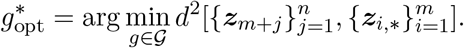

This discrete optimal translation strategy gives both the optimal perturbation representation 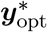 to achieve the desired translation, and also the optimal perturbation 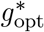. If the cells in the target latent space are treated by a perturbation, we can evaluate if the optimal perturbation 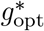 matches the one for the target latent space.

### Integrated gradients

As we connect perturbation and cell state in PerturbNet, we can interpret how a perturbation changes the cell state distribution by predicting cellular representations using PerturbNet. We can further interpret the effects of features and components of the perturbation with the state-of-the-art XAI methods. Denote *F*(·) as a function taking input feature vector ***T*** = (*T*_1_,…,*T_n_*)^*T*^ ∈ ℝ^*n*^ to generate output in [0,1]. Then its attribution is a vector ***A*** = (*α*_1_,…, *a_n_*)^*T*^ and each value *a_i_* is the contribution of *T_i_* to the prediction of *F*(***T***).

Previous attempts to interpret neural network models have focused on gradients [57, 58] and back-propagation [59, 60]. We use the method of integrated gradients [28], which has been applied to interpret deep learning models across a range of domains, including computational chemistry [61]. The attribution score of the integrated gradients method for the *i*th dimension of input ***T*** is defined as

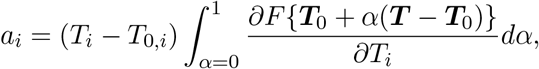

where ***T***_0_ = (*T*_0,0_,…, *T*_0,*n*_)^*T*^ is a baseline input.

A prediction neural network model on cellular representation can be formulated from PerturbNet as ***Z*** = *f*(***V***|***Y***) and ***Y*** = *h*(***B***). The input ***T*** can be formulated as (***V**^T^*, ***Y**^T^*)^*T*^ or (***V**^T^*, ***B**^T^*)^*T*^. In addition, a classification neural network model on ***Z*** provides a classification score within [0, 1]. We can then find input features that increase the probability of generating cells in a particular cell state.

## A Supplementary information

### A.1 Covariate adjustment gives better predictions for PerturbNet

Because the sci-Plex dataset has two covariates (cell type and dose), we adjusted these covariates in modeling cINN translations of PerturbNet. We converted cell type and dose to one-hot encodings and concatenated them to the perturbation representation ***Y*** as a joint condition representation. Then we trained cINN with the joint representation of perturbation and covariates as conditions for translations between residual representation and cellular representation. We then predicted single-cell responses to a perturbation with the specific values of covariates.

We evaluated the prediction performance of PerturbNet adjusted for covariates on the unseen and observed perturbations with cell covariates’ values, and compared its performance with that of the previous PerturbNet trained without the cell state covariates. As can be seen in Supplementary Fig. 1a, the PerturbNet adjusted for cell state covariates significantly outperforms the PerturbNet without covariate adjustment for observed perturbations in both R squared and FID. The PerturbNet adjusted for covariates improves R squared for the unseen perturbations. The cell state covariates are correlated with perturbation assignment and also influence cellular responses, making them possess confounding effects in modeling perturbation responses. Therefore, adjusting for covariates in cINN modeling of the PerturbNet helps debias their confounding effects and more accurately quantify perturbation effects.

We compared the performance of PerturbNet adjusted for covariates with the baseline models. As the PerturbNet adjusted for covariates takes additional covariate information other than perturbation, we performed a stratified prediction in each cell type by dose stratum to also adjust covariate information for the baseline models. Each perturbation has 12 strata with three cell types and four doses. We proceeded with the sampling procedures of the baseline KNN and random models within each cell type by dose stratum, and made PerturbNet predictions with the corresponding covariates’ values in the stratum. Supplementary Fig. 1b-e show that PerturbNet consistently outperforms the random model for observed perturbations, while KNN is unable to beat the random model for either unseen or observed perturbations. As the stratified evaluations constrain cellular variability and sample size, which possibly narrows down the prediction performances of the KNN and random models, we also compared PerturbNet adjusted for covariates and KNN in stratified predictions (Supplementary Fig. 1f-g). As with their unstratified comparisons, the PerturbNet has a better performance for observed perturbations but does not defeat KNN for unseen perturbations.

### A.2 Fine-tuned ChemicalVAE and GenotypeVAE improve the performance of PerturbNet

We performed ChemicalVAE and GenotypeVAE fine-tuning to improve the performance of PerturbNet. To construct a cell-state Laplacian matrix ***L***, we computed the Wasserstein-2 (W2) distance between cellular latent values of each pair of perturbations. As the number of perturbation pairs is extremely large in LINCS-Drug or LINCS-Gene, we first fitted a KNN algorithm on the perturbation representations of a dataset and selected the 30 nearest neighbors for each perturbation to compute their pairwise cellular latent distances. As the resulting pairwise cell latent distance matrix for all the perturbations was not symmetric, we took the average of the matrix and its transpose. We then calculated the exponential of their opposite values and row-normalized the matrix to obtain the adjacency matrix with each entry as a transition probability. We then obtained the Laplacian matrix from the adjacency matrix.

We utilized the Laplacian sub-matrix for the observed chemical perturbations of LINCS-Drug to fine-tune ChemicalVAE, and also that for the observed genetic perturbations of LINCS-Gene to fine-tune GenotypeVAE. We considered values of λ in (0.1, 1, 5, 10, 100, 1000, 10,000) to implement the ChemicalVAE and GenotypeVAE fine-tuning algorithm. After we fine-tuned ChemicalVAE and GenotypeVAE, we evaluated the KNN model on their perturbation representations. We also constructed the cINN model of the PerturbNet between the perturbation representations of the fined-tuned models and cellular representations using cells with the observed perturbations. We evaluated the prediction performance of the fine-tuned KNN and PerturbNet models on the 2000 unseen perturbations of the LINCS-Drug data (Supplementary Fig. 4a). Both R squared and FID of PerturbNet have small to medium fluctuations across increasing λ values, while those of KNN do not obviously change with varying λ values. Several λ values give slight increases of median R squared or decreases of median FID for PerturbNet over the non-fine-tuned one, such as λ = 0.1, 1, 5, 10, 100.

We compared the fine-tuned KNN and PerturbNet with λ = 1 to their non-fine-tuned counterparts for the unseen perturbations of LINCS-Drug (Supplementary Fig. 6b). The fine-tuned PerturbNet has significant improvements in both R squared and FID, while fine-tuning ChemicalVAE does not significantly enhance KNN. A possible explanation is that the cINN of PerturbNet further enforces the prediction capacity from fine-tuned perturbation representation to cell state.

Supplementary Fig. 6c-d show the R squared and FID of KNN and PerturbNet trained with fine-tuned GenotypeVAE (λ = 0.1, 1, 5, 10, 100, 1000, 10000). By comparing the evaluation metrics obtained from fine-tuned KNN and PerturbNet with different λ values, we determined that λ = 1 was the optimal hyperparameter. Supplementary Figure 6e-f shows the scatter plots of R squared and FID of KNN and PerturbNet with fine-tuned GenotypeVAE of λ = 1 over those with non-fine-tuned GenotypeVAE. Fine-tuning GenotypeVAE significantly improves the performance of PerturbNet, especially for observed perturbations. Somewhat surprisingly, the fine-tuning algorithm improves only the PerturbNet, but does not significantly improve the performance of the KNN model.

### A.3 Supplementary figures and tables

**Supplementary Fig. 1.**
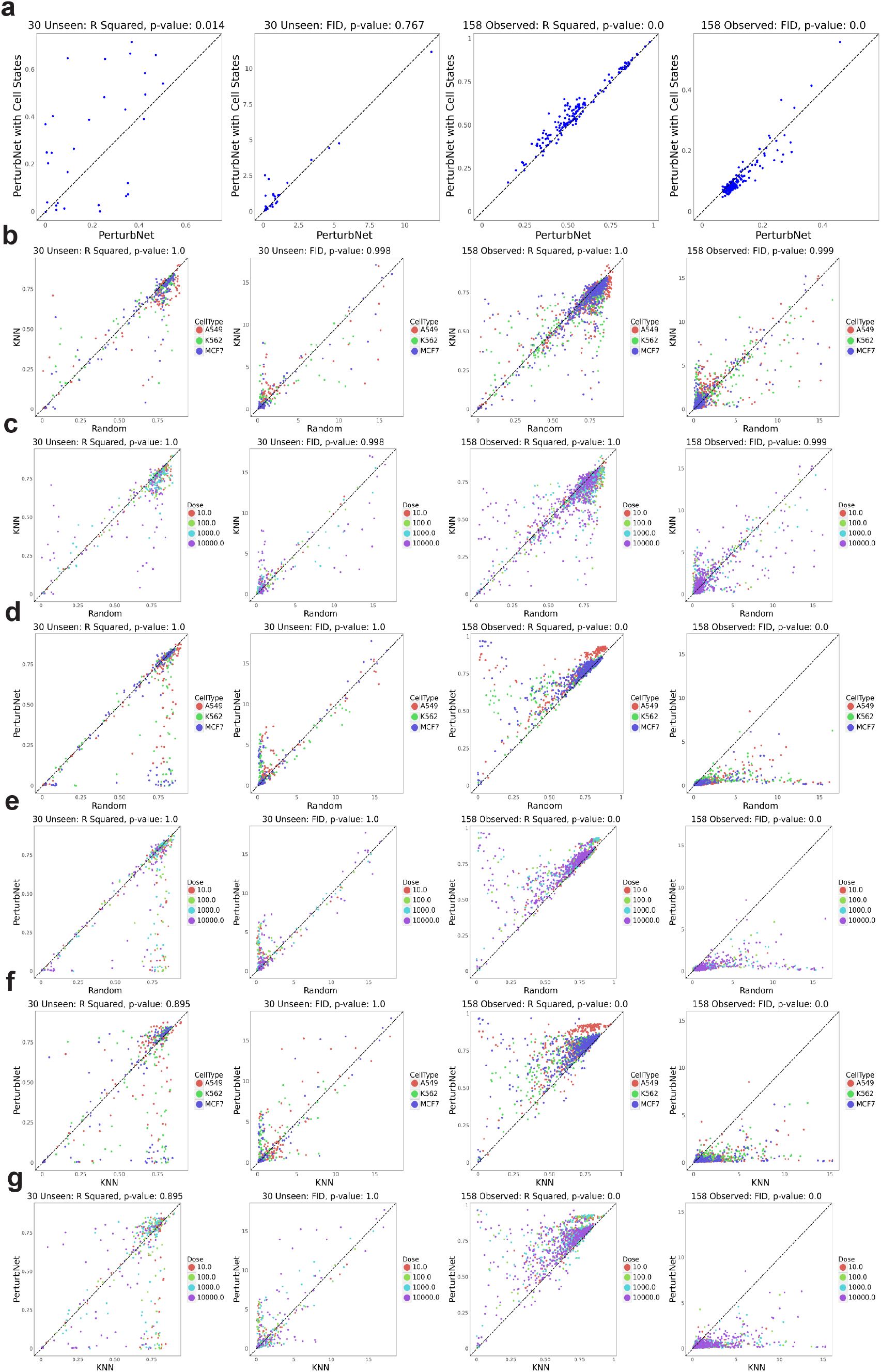
Metrics for PerturbNet with covariate adjustment on sci-Plex dataset. **(a)** Overall comparison between adjusted (y-axis) and unadjusted models. **(b)-(g)** Stratified comparisons among adjusted PerturbNet, KNN, and random model colored by various covariates.

**Supplementary Fig. 2.**
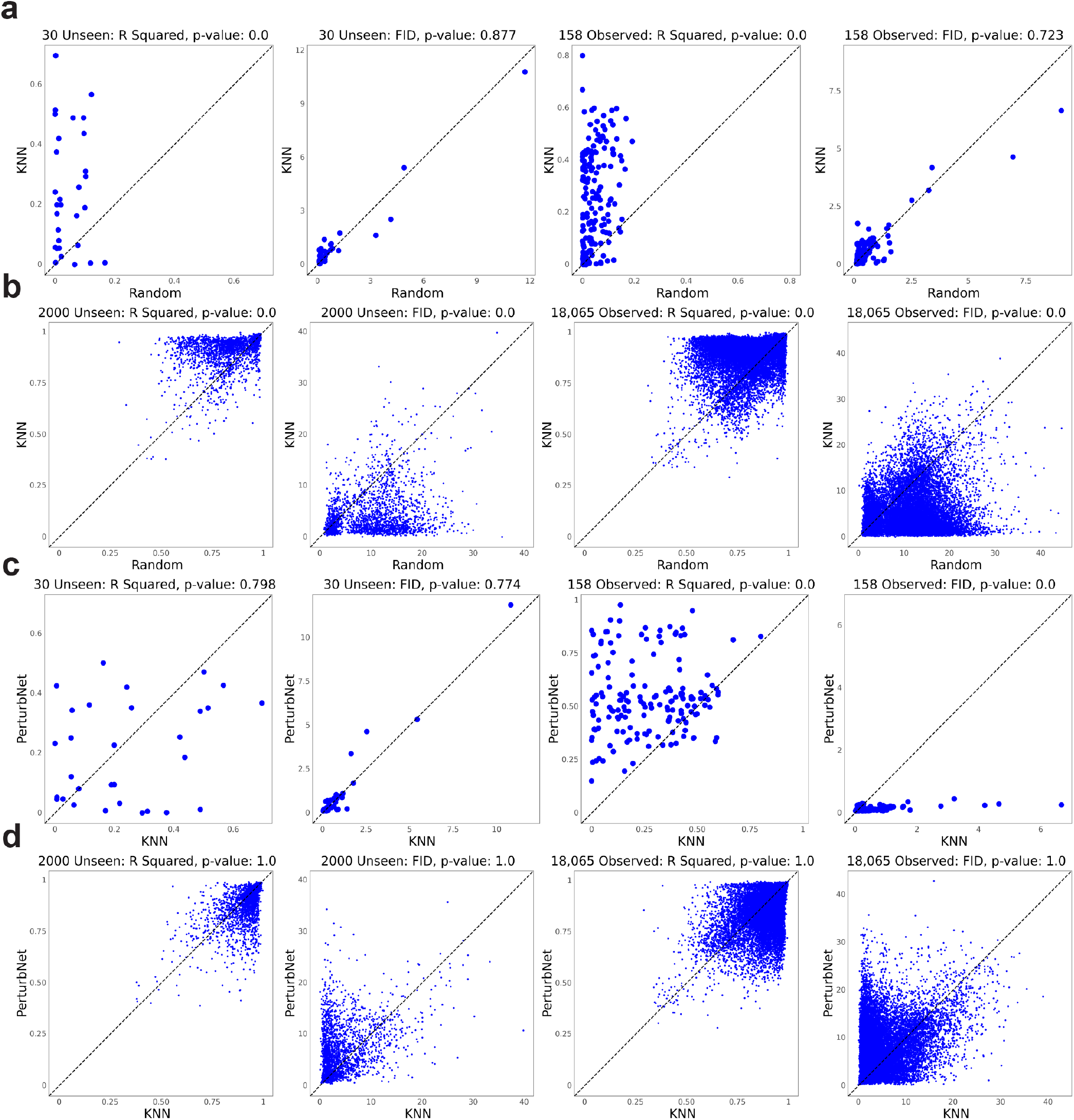
Metrics for KNN method on drug perturbation datasets. **(a)** Unseen and observed sci-Plex perturbations. **(b)** Unseen and observed LINCS-Drug perturbations. **(c)** Unseen and observed sci-Plex perturbations. **(d)** Unseen and observed LINCS-Drug perturbations.

**Supplementary Fig. 3.**
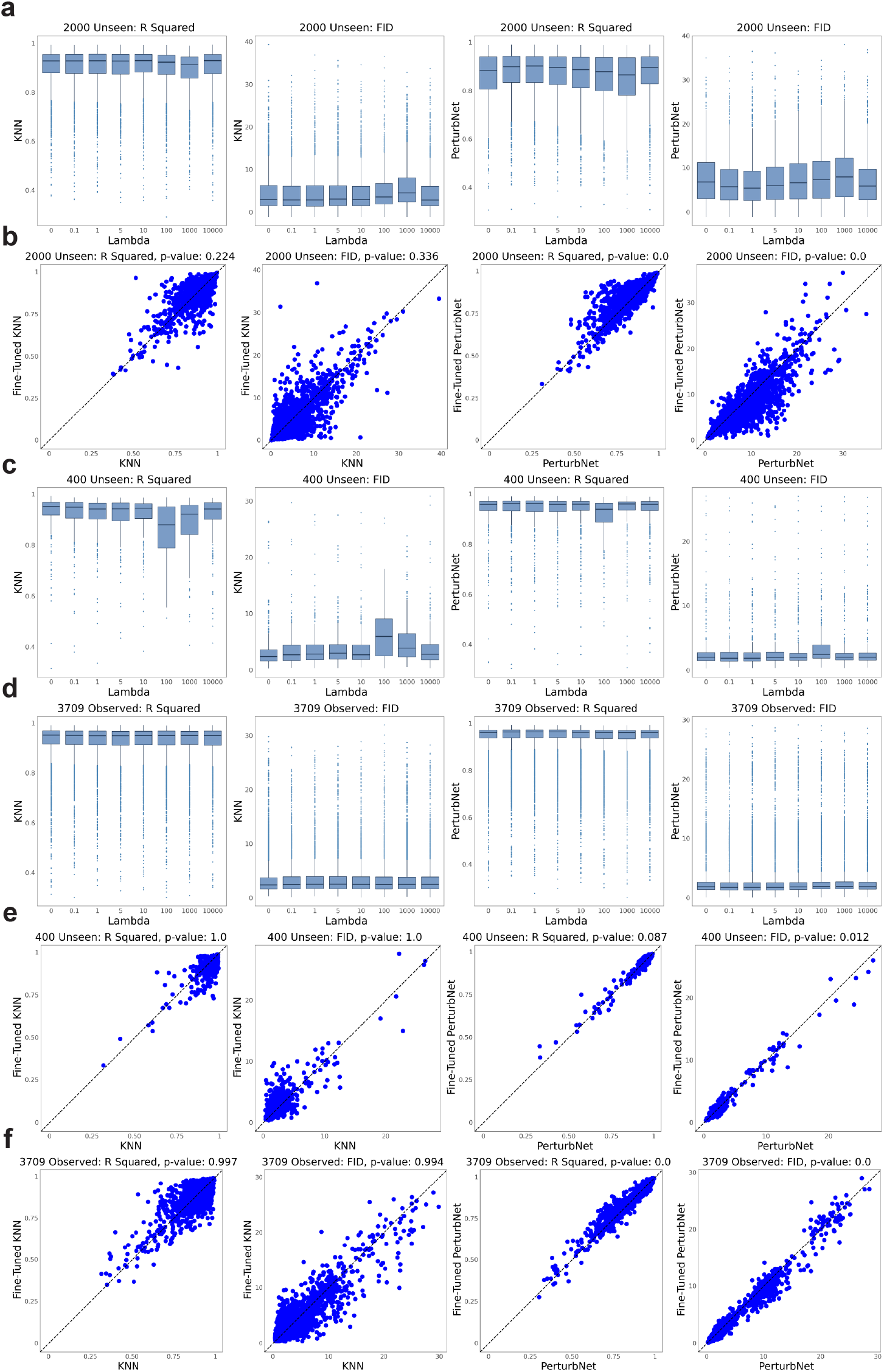
Metrics using fine-tuned perturbation representation networks on LINCS-Drug and LINCS-Gene datasets. **(a)** Box plots of R squared and FID metrics for KNN and PerturbNet on LINCS-Drug (unseen perturbations) after fine-tuning. Lambda is the tuning parameter that balances the ELBO and the graph regularization term. **(b)** Metrics for LINCS-Drug after fine-tuning (λ = 1). **(c)** Box plots of metrics for LINCS-Gene unseen perturbations after fine-tuning with variable λ. **(d)** Box plots of metrics for LINCS-Gene observed perturbations after fine-tuning with variable λ. **(e)-(f)** Metrics for LINCS-Gene unseen (e) and observed (f) perturbations after fine-tuning (λ = 1).

**Supplementary Fig. 4.**
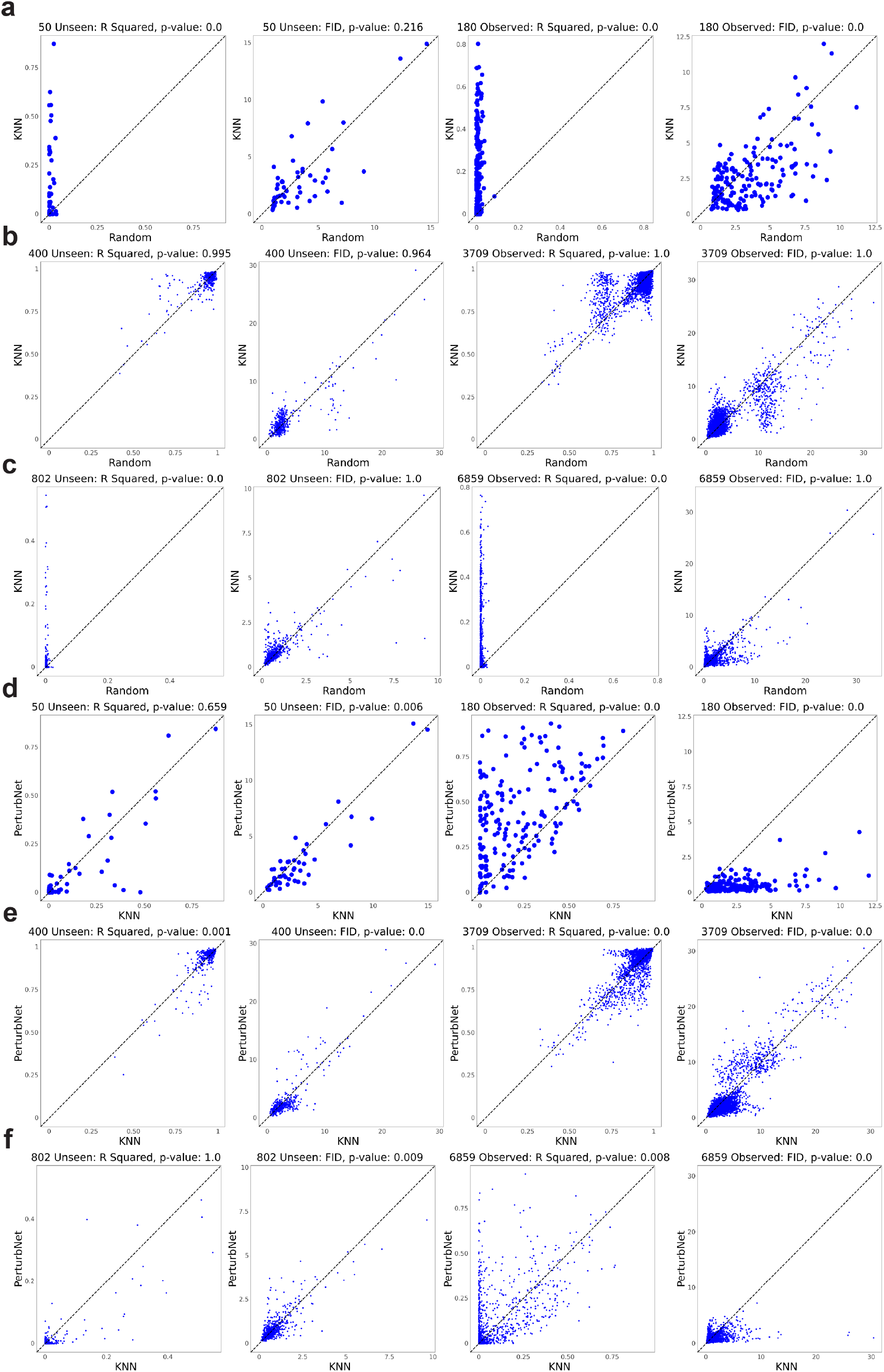
Metrics for KNN method on genetic perturbation datasets. **(a)** Unseen and observed GI perturbations. **(b)** Unseen and observed LINCS-Gene perturbations. **(c)** Unseen and observed GSPS perturbations. **(d)** Unseen and observed GI perturbations. **(e)** Unseen and observed LINCS-Gene perturbations. **(f)** Unseen and observed GSPS perturbations.

**Supplementary Fig. 5.**
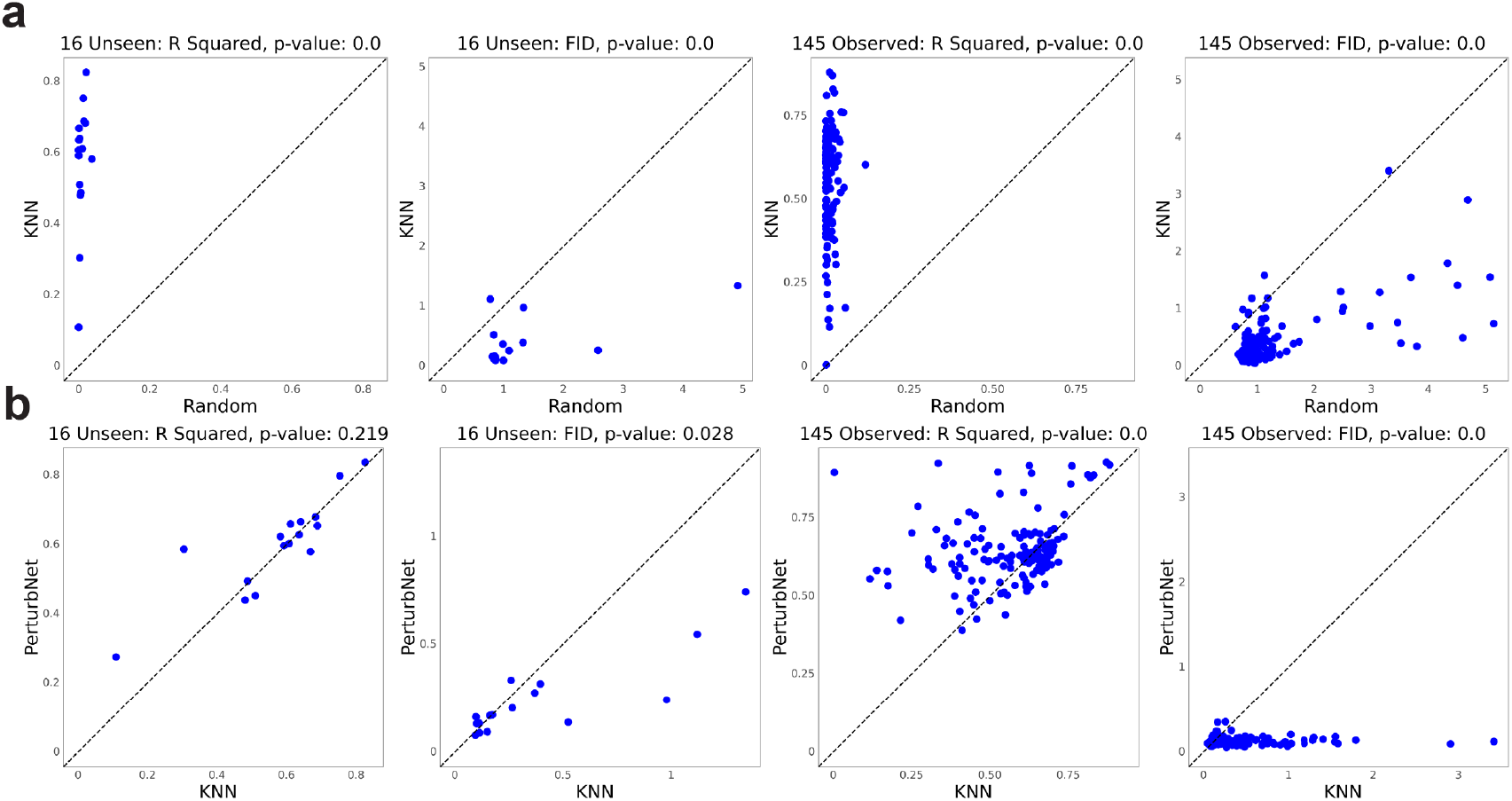
Metrics for KNN method on Ursu dataset. **(a)** KNN vs. random. **(b)** PerturbNet vs. KNN.

**Supplementary Fig. 6.**
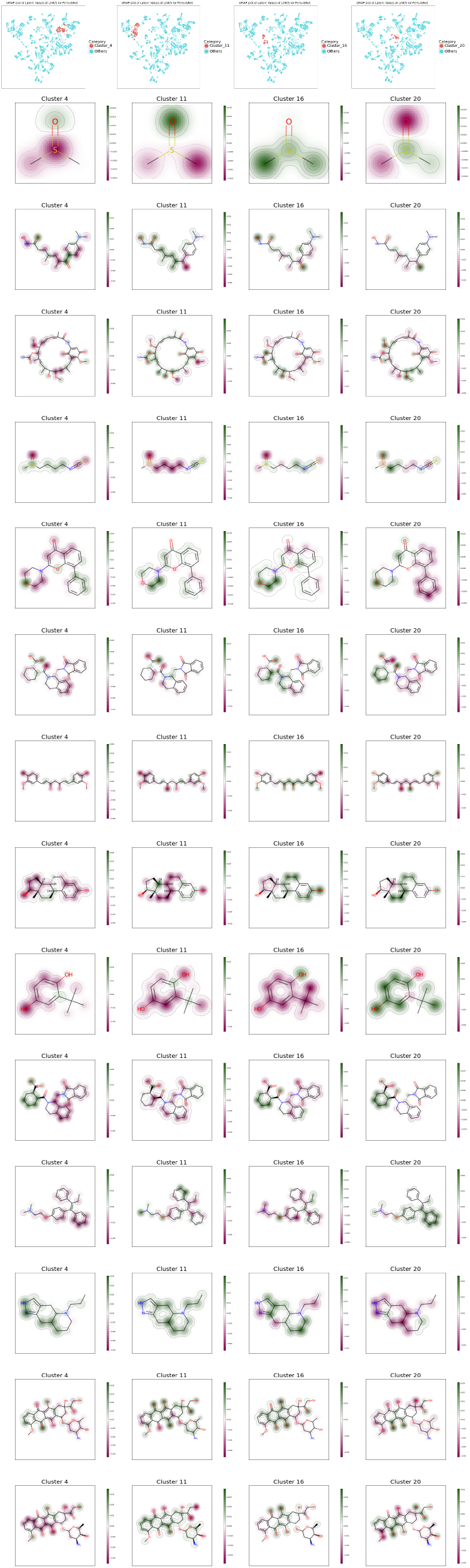
Additional attribution examples from LINCS-Drug dataset. Each column is a different cluster and each row is a different drug.

**Supplementary Fig. 7.**
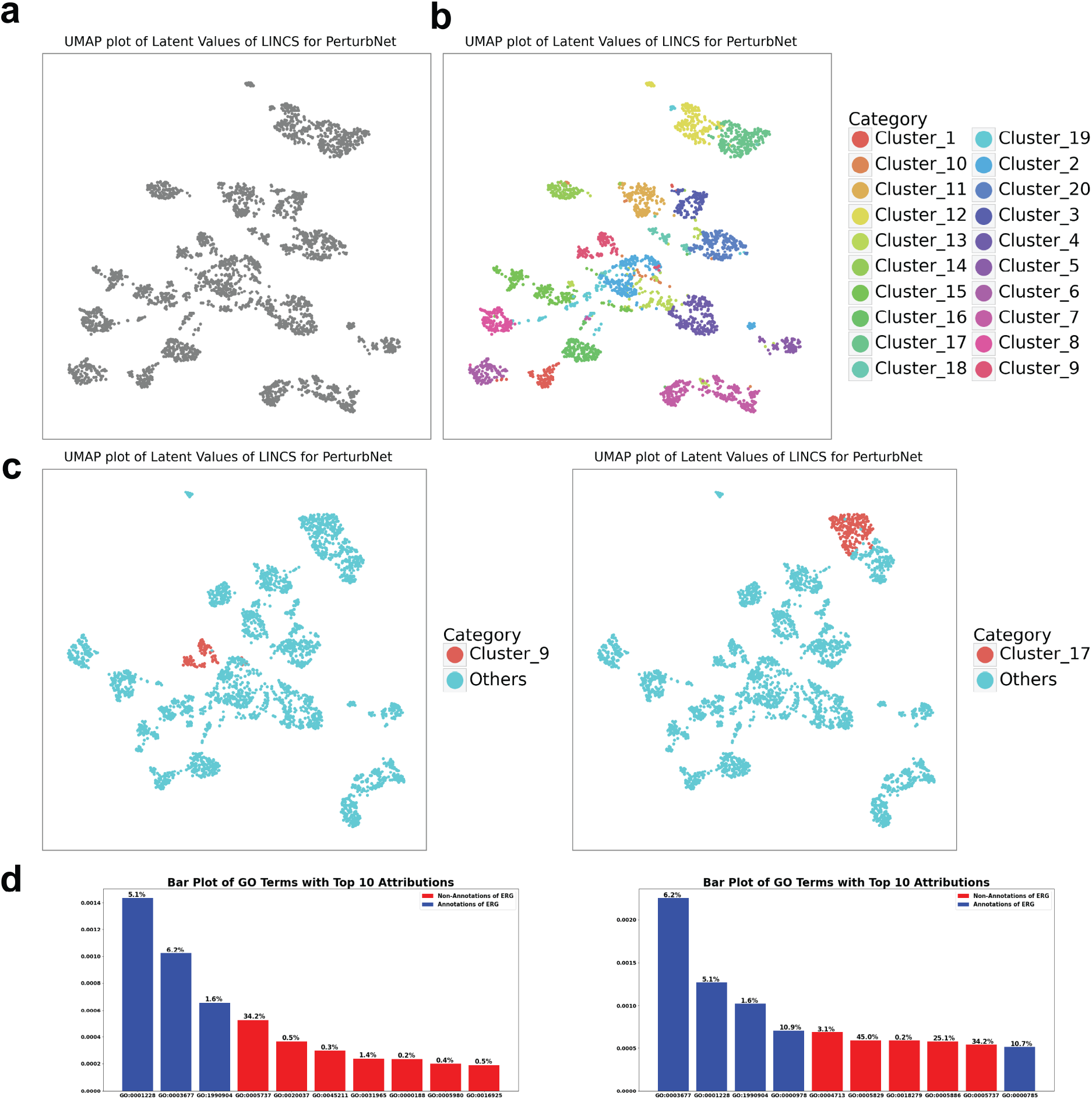
Additional attribution examples from LINCS-Gene dataset. **(a)-(b)** UMAP plots of LINCS-Gene colored gray and by cluster. **(c)** UMAP plots indicating cluster 9 and cluster 17. **(d)** The attribution procedure can give scores for all GO terms, including terms that do not annotate the perturbation of interest. These bar plots show the terms with top 10 attributions, colored by whether they annotate the selected perturbation (ERG). The frequency of the term across all genes is indicated above the bar. Left plot is for cluster 9, right is for cluster 17.

**Supplementary Fig. 8.**
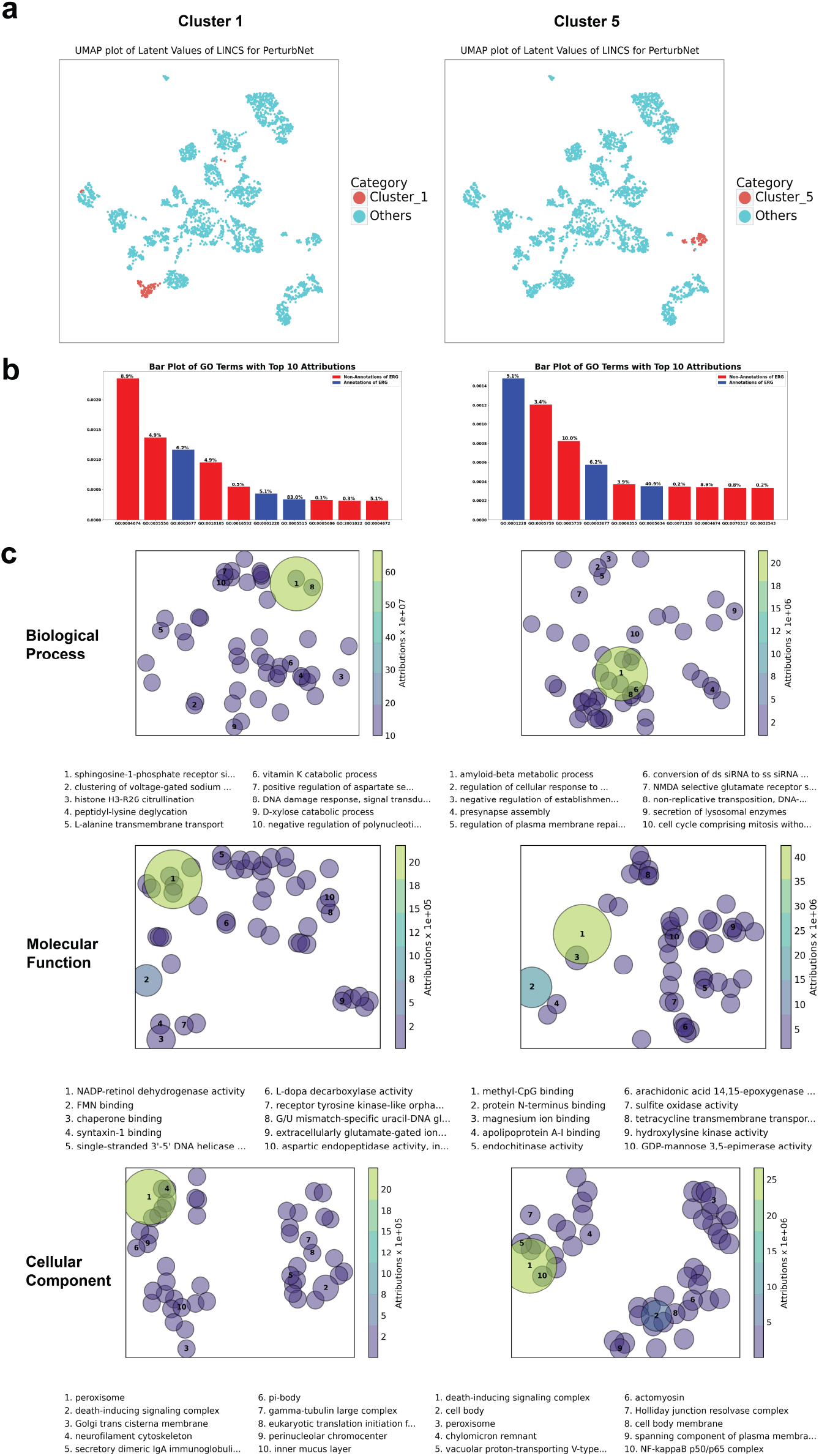
Additional attribution examples from LINCS-Gene dataset. **(a)** UMAP plots indicating cluster 1 and cluster 5. **(b)** Bar plots showing the terms with top 10 attributions, colored by whether they annotate the selected perturbation (ERG). The frequency of the term across all genes is indicated above the bar. Left plot is for cluster 1, right is for cluster 5. **(c)** Plots of GO terms with top attributions for clusters 1 and 5.

## References

[1] Gehring, J., Hwee Park, J., Chen, S., Thomson, M. & Pachter, L. Highly multiplexed single-cell rna-seq by dna oligonucleotide tagging of cellular proteins. Nature Biotechnology 38, 35–38 (2020).

[2] Srivatsan, S. R. et al. Massively multiplex chemical transcriptomics at single-cell resolution. Science 367, 45–51 (2020).

[3] Doudna, J. A. & Charpentier, E. The new frontier of genome engineering with crispr-cas9. Science 346(2014).

[4] Dixit, A. et al. Perturb-seq: dissecting molecular circuits with scalable single-cell rna profiling of pooled genetic screens. cell 167, 1853–1866 (2016).

[5] Wang, T. et al. Identification and characterization of essential genes in the human genome. Science 350, 1096–1101 (2015).

[6] Adamson, B. et al. A multiplexed single-cell crispr screening platform enables systematic dissection of the unfolded protein response. Cell 167, 1867–1882 (2016).

[7] Datlinger, P. et al. Pooled crispr screening with single-cell transcriptome readout. Nature methods 14, 297–301 (2017).

[8] Ursu, O. et al. Massively parallel phenotyping of variant impact in cancer with perturb-seq reveals a shift in the spectrum of cell states induced by somatic mutations. bioRxiv (2020).

[9] Jin, X. et al. In vivo perturb-seq reveals neuronal and glial abnormalities associated with autism risk genes. Science 370(2020).

[10] Norman, T. M. et al. Exploring genetic interaction manifolds constructed from rich single-cell phenotypes. Science 365, 786–793 (2019).

[11] Lotfollahi, M., Wolf, F. A. & Theis, F. J. scgen predicts single-cell perturbation responses. Nature methods 16, 715–721 (2019).

[12] Rampáŝek, L., Hidru, D., Smirnov, P., Haibe-Kains, B. & Goldenberg, A. Dr. vae: improving drug response prediction via modeling of drug perturbation effects. Bioinformatics 35, 3743–3751 (2019).

[13] Lotfollahi, M., Naghipourfar, M., Theis, F. J. &Wolf, F. A. Conditional out-of-distribution generation for unpaired data using transfer vae. Bioinformatics 36, i610–i617 (2020).

[14] Johansson, F., Shalit, U. & Sontag, D. Learning representations for counterfactual inference. In International conference on machine learning, 3020–3029 (PMLR, 2016).

[15] Burkhardt, D. B. et al. Quantifying the effect of experimental perturbations at single-cell resolution. Nature biotechnology 39, 619–629 (2021).

[16] Lotfollahi, M. et al. Compositional perturbation autoencoder for single-cell response modeling. bioRxiv (2021).

[17] Yeo, G. H. T., Saksena, S. D. & Gifford, D. K. Generative modeling of single-cell time series with prescient enables prediction of cell trajectories with interventions. Nature communications 12, 1–12 (2021).

[18] Kamimoto, K., Hoffmann, C. M. & Morris, S. A. Celloracle: Dissecting cell identity via network inference and in silico gene perturbation. bioRxiv (2020).

[19] Papamakarios, G., Nalisnick, E., Rezende, D. J., Mohamed, S. & Lakshminarayanan, B. Normalizing flows for probabilistic modeling and inference. Journal of Machine Learning Research 22, 1–64 (2021).

[20] Baltruŝsaitis, T., Ahuja, C. & Morency, L.-P. Multimodal machine learning: A survey and taxonomy. IEEE transactions on pattern analysis and machine intelligence 41, 423–443 (2018).

[21] Rombach, R., Esser, P. & Ommer, B. Network-to-network translation with conditional invertible neural networks. arXiv preprint arXiv:2005.13580 (2020).

[22] Irwin, J. J. & Shoichet, B. K. Zinc-a free database of commercially available compounds for virtual screening. Journal of chemical information and modeling 45, 177–182 (2005).

[23] Subramanian, A. et al. A next generation connectivity map: L1000 platform and the first 1,000,000 profiles. Cell 171, 1437–1452 (2017).

[24] Chicco, D., Sadowski, P. & Baldi, P. Deep autoencoder neural networks for gene ontology annotation predictions. In Proceedings of the 5th ACM conference on bioinformatics, computational biology, and health informatics, 533–540 (2014).

[25] Replogle, J. M. et al. Mapping information-rich genotype-phenotype landscapes with genome-scale perturb-seq. Cell (2022).

[26] Lopez, R., Regier, J., Cole, M. B., Jordan, M. I. & Yosef, N. Deep generative modeling for single-cell transcriptomics. Nature methods 15, 1053–1058 (2018).

[27] Rives, A. et al. Biological structure and function emerge from scaling unsupervised learning to 250 million protein sequences. Proceedings of the National Academy of Sciences 118(2021).

[28] Sundararajan, M., Taly, A. & Yan, Q. Axiomatic attribution for deep networks. In International conference on machine learning, 3319–3328 (PMLR, 2017).

[29] Schiebinger, G. et al. Optimal-transport analysis of single-cell gene expression identifies developmental trajectories in reprogramming. Cell 176, 928–943 (2019).

[30] Crowley, L. et al. A single-cell atlas of the mouse and human prostate reveals heterogeneity and conservation of epithelial progenitors. Elife 9, e59465 (2020).

[31] Demetci, P., Santorella, R., Sandstede, B., Noble, W. S. & Singh, R. Gromov-wasserstein optimal transport to align single-cell multi-omics data. BioRxiv (2020).

[32] Bergen, V., Lange, M., Peidli, S., Wolf, F. A. & Theis, F. J. Generalizing rna velocity to transient cell states through dynamical modeling. Nature biotechnology 38, 1408–1414 (2020).

[33] Kim, S. et al. Pubchem substance and compound databases. Nucleic acids research 44, D1202–D1213 (2016).

[34] Lotfollahi, M. et al. Mapping single-cell data to reference atlases by transfer learning. Nature Biotechnology 40, 121–130 (2022).

[35] Yu, H. & Welch, J. D. Michigan: sampling from disentangled representations of single-cell data using generative adversarial networks. Genome biology 22, 1–26 (2021).

[36] Landrum, G. Rdkit: open-source cheminformatics http://www.rdkit.org. Google Scholar There is no corresponding record for this reference (2016).

[37] Wolf, F. A., Angerer, P. & Theis, F. J. Scanpy: large-scale single-cell gene expression data analysis. Genome biology 19, 15 (2018).

[38] Rogers, D. & Hahn, M. Extended-connectivity fingerprints. Journal of chemical information and modeling 50, 742–754 (2010).

[39] Xu, Z., Wang, S., Zhu, F. & Huang, J. Seq2seq fingerprint: An unsupervised deep molecular embedding for drug discovery. In Proceedings of the 8th ACM international conference on bioinformatics, computational biology, and health informatics, 285–294 (2017).

[40] Chithrananda, S., Grand, G. & Ramsundar, B. Chemberta: Large-scale self-supervised pretraining for molecular property prediction. arXiv preprint arXiv:2010.09885 (2020).

[41] Kusner, M. J., Paige, B. & Hernández-Lobato, J. M. Grammar variational autoencoder. In International Conference on Machine Learning, 1945–1954 (PMLR, 2017).

[42] Zhu, J. et al. Prediction of drug efficacy from transcriptional profiles with deep learning. Nature Biotechnology 1–9 (2021).

[43] Gómez-Bombarelli, R. et al. Automatic chemical design using a data-driven continuous representation of molecules. ACS central science 4, 268–276 (2018).

[44] Ma, J. et al. Using deep learning to model the hierarchical structure and function of a cell. Nature methods 15, 290–298 (2018).

[45] Devlin, J., Chang, M.-W., Lee, K. & Toutanova, K. Bert: Pre-training of deep bidirectional transformers for language understanding. arXiv preprint arXiv:1810.04805 (2018).

[46] Vaswani, A. et al. Attention is all you need. Advances in neural information processing systems 30(2017).

[47] Rao, R. M. et al. Msa transformer. In International Conference on Machine Learning, 8844–8856 (PMLR, 2021).

[48] Meier, J. et al. Language models enable zero-shot prediction of the effects of mutations on protein function. Advances in Neural Information Processing Systems 34(2021).

[49] Heusel, M., Ramsauer, H., Unterthiner, T., Nessler, B. & Hochreiter, S. Gans trained by a two time-scale update rule converge to a local nash equilibrium. In Advances in neural information processing systems, 6626–6637 (2017).

[50] Dinh, L., Sohl-Dickstein, J. & Bengio, S. Density estimation using real nvp. arXiv preprint arXiv:1605.08803 (2016).

[51] Ardizzone, L., Lüth, C., Kruse, J., Rother, C. & Köthe, U. Guided image generation with conditional invertible neural networks. arXiv preprint arXiv:1907.02392 (2019).

[52] Kingma, D. P. & Dhariwal, P. Glow: Generative flow with invertible 1×1 convolutions. Advances in neural information processing systems 31(2018).

[53] Cai, D., He, X., Han, J. & Huang, T. S. Graph regularized nonnegative matrix factorization for data representation. IEEE transactions on pattern analysis and machine intelligence 33, 1548–1560 (2010).

[54] Kingma, D. P. & Ba, J. Adam: A method for stochastic optimization. arXiv preprint airXiv:1412.6980 (2014).

[55] Vaserstein, L. N. Markov processes over denumerable products of spaces, describing large systems of automata. Problemy Peredachi Informatsii 5, 64–72 (1969).

[56] Dowson, D. & Landau, B. The fréchet distance between multivariate normal distributions. Journal of multivariate analysis 12, 450–455 (1982).

[57] Baehrens, D. et al. How to explain individual classification decisions. The Journal of Machine Learning Research 11, 1803–1831 (2010).

[58] Simonyan, K., Vedaldi, A. & Zisserman, A. Deep inside convolutional networks: Visualising image classification models and saliency maps. arXiv preprint arXiv:1312.6034 (2013).

[59] Shrikumar, A., Greenside, P., Shcherbina, A. & Kundaje, A. Not just a black box: Learning important features through propagating activation differences. arXiv preprint arXiv:1605.01713 (2016).

[60] Shrikumar, A., Greenside, P. & Kundaje, A. Learning important features through propagating activation differences. In International conference on machine learning, 3145–3153 (PMLR, 2017).

[61] McCloskey, K., Taly, A., Monti, F., Brenner, M. P. & Colwell, L. J. Using attribution to decode binding mechanism in neural network models for chemistry. Proceedings of the National Academy of Sciences 116, 11624–11629 (2019).

